# Phage-encoded sRNA counteracts xenogenic silencing in pathogenic *E. coli*

**DOI:** 10.1101/2025.11.27.690950

**Authors:** Pranita Poudyal, Brandon Sy, Daniel G. Mediati, Michael Payne, Vibhuti Nandel, Dougall Norris, Sean McAteer, Asim Ullah, Serena Li, Lawrence Menz, Shafagh A. Waters, Timothy J. Dallman, Mathew Baker, Ruiting Lan, David Gally, Jai Tree

## Abstract

Horizontal gene transfer introduces foreign DNA that can disrupt cellular processes and is therefore subject to xenogenic silencing by nucleoid-associated proteins such as H-NS and Hha. In Enterohaemorrhagic *Escherichia coli* (EHEC), prophages make up a large fraction of the accessory genome and encode many virulence factors, yet to be expressed they must overcome this silencing. We identify a prophage-encoded small RNA (sRNA), HnrS, that functions as an anti-silencing factor by targeting the H-NS paralogue Hha. HnrS is a short (66-nt) sRNA present in multiple copies (up to nine) in EHEC and Enteropathogenic *Escherichia coli* (EPEC) genomes and is enriched in *E. coli* strains that carry the locus of enterocyte effacement (LEE⁺). We show that HnrS directly base-pairs with the ribosome-binding site of the *hha* mRNA, repressing its translation and thereby reducing Hha-enhanced H-NS silencing. This counter-silencing de-represses the LEE type III secretion system and concomitantly represses motility. Transcriptomic profiling further revealed that HnrS indirectly activates genes involved in nitrate/nitrite respiration and nitric oxide resistance, metabolic pathways that contribute to survival in the inflamed gastrointestinal tract. Deletion of *hnrS* reduced expression of nitrate reductase genes and impaired actin pedestal formation on host epithelial cells. Our results indicate that prophage-encoded, multicopy *hnrS* provides a counter-silencing mechanism that reduces Hha–H-NS repression at specific virulence loci. This likely enables expression of horizontally acquired genes without broadly disrupting the core H-NS regulon. HnrS illustrates how mobile genetic elements deploy sRNAs to counteract xenogenic silencing and promote virulence gene expression, enhancing colonisation of the host.

**Importance statement:** Horizontally acquired genes are often silenced to protect bacterial genomes, but this defence also limits the expression of new traits. We identify a prophage-encoded small RNA, HnrS, that counteracts this restriction by repressing the xenogenic silencer Hha, lifting Hha–H-NS-mediated repression of virulence and metabolic genes. HnrS activates the locus of enterocyte effacement, nitrate/nitrite respiration, and nitric oxide resistance—pathways that help *E. coli* survive and colonise the inflamed gut. Our findings reveal an RNA-based counter-silencing mechanism encoded by mobile genetic elements, showing how phages can reprogram bacterial regulatory networks to promote adaptation and pathogenicity.

## INTRODUCTION

Horizontal gene transfer (HGT) plays a central role in bacterial evolution, enabling the acquisition of genes that confer new functions such as antibiotic resistance and virulence. However, incoming DNA can also disrupt cellular processes or impose metabolic costs and is therefore restricted by multiple layers of genome defence. The most frequent include restriction–modification systems, CRISPR-Cas, and cyclic oligonucleotide-based antiphage signalling systems (CBASS), among an expanding repertoire of defence systems (1).

Foreign DNA that evades initial restriction can be integrated into the genome but is frequently detrimental to host fitness (2). In *Enterobacteriaceae*, foreign DNA is often more AT-rich than the host genome and is transcriptionally silenced by the nucleoid-associated protein H-NS, which binds to AT-rich regions and forms oligomeric complexes that repress transcription (3–8). This process—termed xenogenic silencing—enables bacteria to tolerate the presence of foreign genes without incurring their deleterious effects (2, 9–11). Deletion of *hns* is lethal in several pathogens (12–15) underscoring the role of H-NS in suppressing deleterious gene expression.

For horizontally-acquired genes to be retained under positive selection they must be expressed. Activation can occur through specific transcription factors that counteract H-NS-mediated silencing in response to environmental cues (15). A broader anti-silencing strategy is used by bacteriophage T7 that encodes a protein (gene 5.5) that forms heterodimers with H-NS, reducing its DNA-binding activity and enabling phage transcription (16). However, global interference with H-NS is likely to de-repress a broad range of largely detrimental genes and be costly to the host.

H-NS silencing is selectively enhanced at virulence loci by Hha, a small H-NS paralogue that lacks a DNA-binding domain but forms heterodimers with H-NS to increase its affinity for specific targets (17). Hha appears to enhance H-NS activity on foreign genes, particularly at virulence-associated islands, without affecting the expression of H-NS-regulated genes in the core genome (17, 18). Structural and biochemical analysis of the Hha-H-NS heterodimer indicates that Hha coats the H-NS oligomerisation domain and presents a positive surface that increases DNA binding (18).

In Enterohaemorrhagic *Escherichia coli* (EHEC), horizontal gene transfer has introduced over 1.5 Mb of new genetic material, including 19 prophages that carry many key virulence factors (19, 20). We previously showed that these regions are rich in small regulatory RNAs (sRNAs), several of which contribute to virulence gene expression (21–23). Here, we identify an sRNA, HnrS, encoded on prophages and present in up to nine copies in attaching and effacing *E. coli* strains that encode the locus of enterocyte effacement (LEE+). We demonstrate that HnrS directly base pairs with the ribosome binding site of *hha* mRNA to represses *hha* expression, partly relieving Hha-H-NS-mediated silencing. This results in upregulation of the LEE encoded type 3 secretion system and repression of motility. In addition, HnrS promotes the expression of genes involved in nitric oxide resistance and nitrate/nitrite respiration—key adaptations for survival in the inflamed gastrointestinal tract (24–26). Together, these data support a model in which multiple prophage-encoded copies of HnrS provide a dosage-tunable mechanism to counteract xenogenic silencing at specific virulence loci, promoting virulence gene expression and host colonisation while avoiding broad de-repression of the core H-NS regulon.

## RESULTS

### HnrS is a multicopy small RNA encoded on prophages

In EHEC strains horizontally-acquired pathogenicity islands encode the majority of virulence factors, as well as regulators that tune their expression. In our previous analysis of the sRNA interactome in EHEC str. Sakai we identified several sRNAs within the pathogenicity islands (S-loops) with abundant mRNA interactions (27), suggesting a role in virulence. Among the most abundant were three identical copies of a single sRNA – designated EcOnc10, EcOnc11, and EcOnc12 – encoded within the prophages Sp9, Sp10, and Sp12 respectively. We here rename these sRNA HnrS (Hha-negative regulatory small RNA). We had previously identified these sRNAs through their Hfq-binding (21) indicating that they are Hfq-associated sRNA (**Figure 1A**). To precisely map the 5’ and 3’ ends of the sRNAs, we analysed dRNA-seq and Term-seq data for EHEC str. Sakai. All three loci share identical 5’ and 3’ ends and produced a 66-nt sRNA transcript terminated by an intrinsic Rho-independent terminator (**Figure 1A**). Northern blot analysis supported transcription of a 66-nt sRNA and indicated that it accumulates during early stationary phase in both T3SS-inducing MEM-HEPES and minimal M9 media (**Figure 1B**), similar to the Stx phage sRNA, StxS (28)

**Figure 1.**
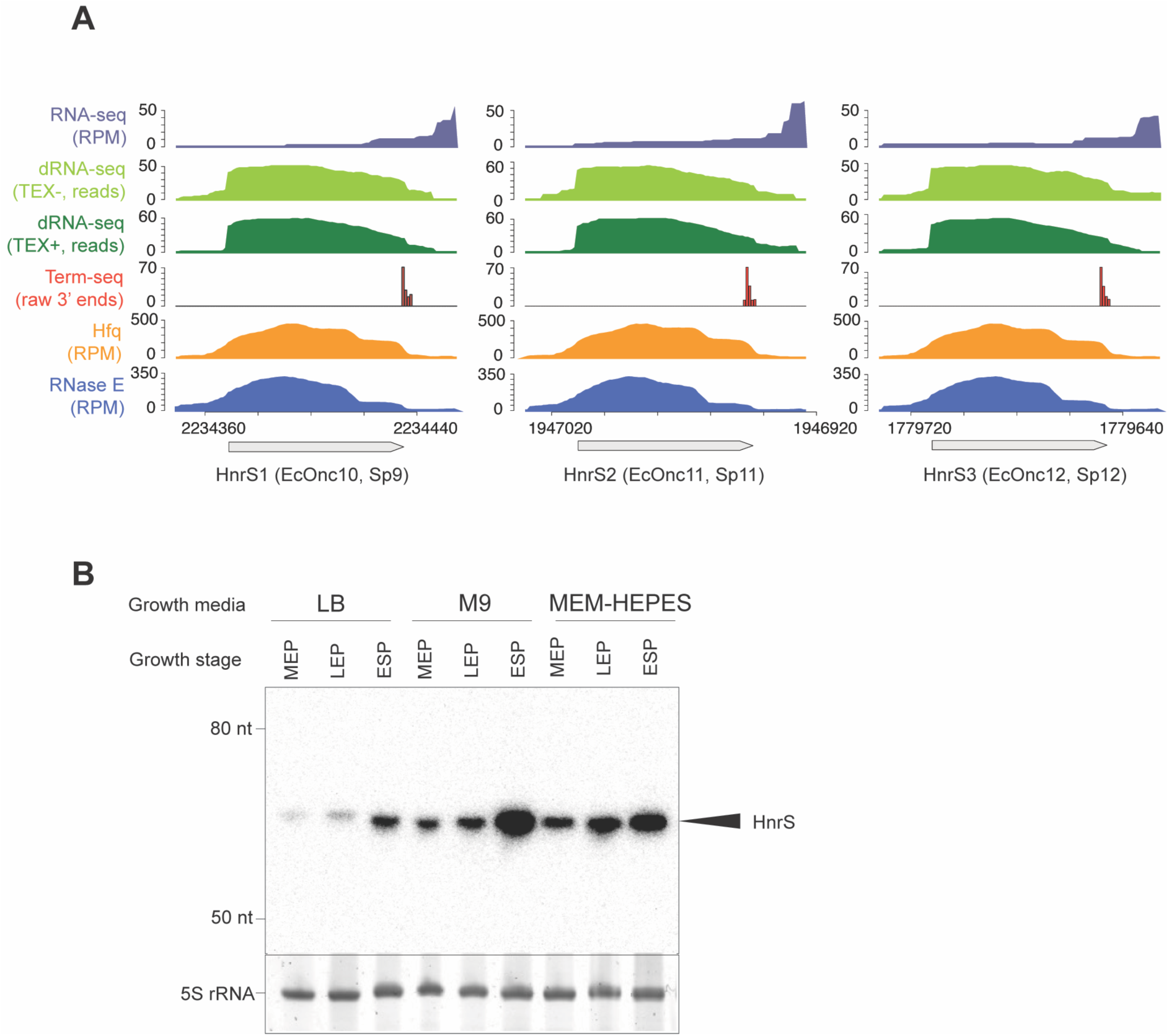
*E. coli* str. Sakai encodes three identical copies of sRNA HnrS on prophage. **A.** RNA end mapping, Hfq, and RNase E binding at HnrS. Total RNA-seq data from EHEC grown in MEM-HEPES (purple, [23]), dRNA-seq (light green, TEX-), dRNA-seq dark green, TEX+), and Term-seq (red bar plot), Hfq binding (orange, [21]), and RNase E binding (dark blue, [22]) for three genomic copies of HnrS in EHEC str. Sakai. Read abundance is indicated on the y-axis. Term-seq peaks correspond to transcription termination sites, while dRNA-seq reads highlight transcription start sites. Grey arrows indicate the position of HnrS with the systematic name and prophages (Sp9, Sp10, and Sp12) in brackets. **B.** Northern blot analysis of HnrS expression in LB, minimal M9, and MEM-HEPES media at mid-exponential (OD600 = 0.6), late exponential (OD600 = 1.0), and early stationary (OD600 = 1.8) phases.

To assess the distribution of HnrS across *E. coli* pathotypes, we searched for copies within 3,230 *E. coli* genomes in NCBI and assigned pathotypes from characteristic virulence genes (Methods & Supplementary Information)(29, 30). We identified *hnrS* predominantly in EHEC and EPEC pathotypes (98.01%), with a minor representation in uropathogenic (0.82%) and unclassified (1.17%) *E. coli* strains (**Figure 2A**). Shiga toxin genes (*stx1* and s*tx 2* that partly characterise EHEC) were significantly enriched in the *hnrS* encoding strains (*p*<0.0001), as was the LEE that facilitates attaching and effacing lesion formation in EHEC and EPEC (*p*<0.0001, Fishers exact test).

**Figure 2.**
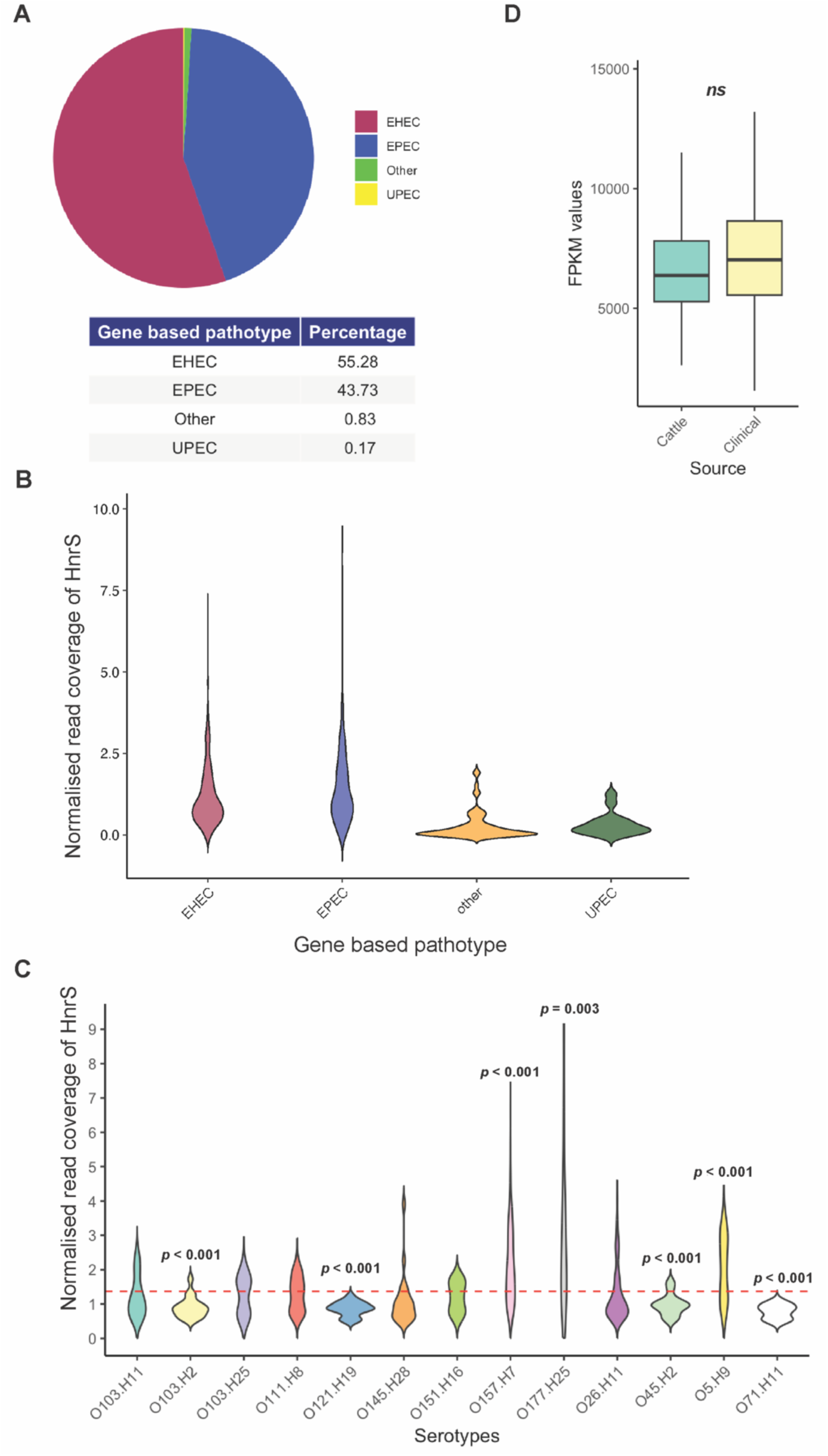
Distribution and coverage of HnrS across *E. coli* pathotypes and serotypes. **A.** Coverage of *hnrS* across different *E. coli* pathotypes, including EHEC, EPEC, UPEC, and others where values were normalised to six conserved housekeeping genes (*arpA*, *trpA*, *uidA*, *rrl*, *tspE4*, and *fyuA*) as a reference. **B.** Violin plot representing maximum *hnrS* coverage across EHEC, EPEC and UPEC strains. **C.** Distribution of *hnrS* read coverage across STEC serotypes. Asterisks above the violins indicate statistically significant deviations from the overall median, as determined by one-sample t-tests. **D.** Box plot of *hnrS* coverage (FPKM) in O157:H7 isolates from human and cattle sources.

The Sakai strain of EHEC has been extensively studied and harbours three copies of HnrS on distinct prophages. To assess variation in gene dosage across strains, we estimated relative *hnrS* copy number from normalised sequencing read depth **(Figure 2A-B**). Specifically, the coverage of *hnrS* was normalised against the mean coverage of six conserved housekeeping genes (*arpA*, *trpA*, *uidA*, *rrl*, *tspE4*, and *fyuA*) to yield a relative read coverage that approximates copy number. The median relative coverage for *hnrS* among O157:H7 strains was approximately 2-fold, with the highest exceeding 7-fold in one isolate **(Figure 2C**).

Among EHEC serotypes, O177:H25 and O157:H7 exhibited a significantly higher *hnrS* relative coverage, with some isolates encoding 9-fold more *hnrS* relative to the housekeeping gene controls. Because O157:H7 is the serotype most commonly associated with human infection, we asked whether increased *hnrS* copy number is associated with human infection. Comparing isolates from human infections versus cattle (reservoir) measured as normalised read depth (FPKM [Fragments per kilobase of transcript per million mapped reads]) (31), indicated that the median relative coverage for human isolates trended higher but this difference was not statistically significant (**Figure 2D**).

Given recent reports of bacteriophage sRNAs that promote phage survival (32, 33), we asked whether HnrS is prophage-encoded in other EHEC and EPEC strains. We used PHASTER analysis to identify prophage regions in a subset of strains. In each EPEC and EHEC strain, a single copy of HnrS was encoded within the late genes of an independent prophage element (**Supplementary Figure 1**). These results indicate that HnrS is acquired by insertion of prophages into the genome, and that multiple copies arise from the acquisition of distinct prophages.

### HnrS represses the xenogenic DNA silencer, Hha

The acquisition of multiple copies of *hnrS* on phage suggests that the sRNA contributes to phage and/or host fitness. Using our previously published sRNA interactome data (RNase E-CLASH) (27), we identified target mRNAs for HnrS. The most abundant interaction for HnrS was with the toxin-antitoxin system, *mazEF* (129-hybrids, FDR=0). The *mazEFG* operon comprises the antitoxin *mazE* and the toxin *mazF*, followed by *mazG*, which encodes a nucleoside triphosphate pyrophosphohydrolase thought to modulate programmed cell death and stress adaptation (34). HnrS has perfect complementary with the last 15-nt the *mazF* coding sequence (**Supplementary Figure 2A)**. We confirmed that HnrS interacts with the *mazF* coding sequence *in vitro* using EMSA (**Supplementary Figure 2B-C**). MazEF has previously been proposed to confer resistance to phage P1 and T4 (34, 35), we therefore tested whether HnrS may antagonise this phage defence system. We did not observe P1 or T4 susceptibility in a Δ*mazEFG* strain and overexpression of HnrS in commensal *E. coli* str. MG1655 (that lacks the *hnrS*-encoding prophages) did not confer phage susceptibility (**Supplementary Figure 2D**). Notably, our strain backgrounds and *ΔmazEFG* construct differ from those previously reported and may explain our divergent results (34, 35). We next asked if HnrS can suppress MazEF toxicity. Overexpression of *mazEF* was toxic to *E. coli* str. MG1655, but co-expression of HnrS had no appreciable effect on MazEF toxicity (**Supplementary Figure 3A**). To understand if HnrS might direct processing of the *mazFG* intergenic region or regulate downstream *mazG*, we constructed a *mazFG*-GFP translational fusion and monitored fluorescence in the presence or absence of HnrS. MazG’-GFP was modestly reduced (∼13%), a change we do not consider biologically meaningful (**Supplementary Figure 3B**). Together these data suggest that while HnrS interacts with *mazF in vivo* and *in vitro*, it does not measurably modulate MazEF abundance or promote T4 or P1 susceptibility in *E. coli* str. MG1655. We note that non-canonical sRNA functions have recently been uncovered (36), leaving open alternative mechanisms not captured by our assays.

To identify functional targets of HnrS, we constructed GFP translational fusions to the remaining HnrS-mRNA interactions identified by RNase E-CLASH (FDR<0.2 or hybrid count>2). Two mRNAs were significantly regulated by HnrS, the transcription factor AsnC was upregulated (1.32-fold, *p*=0.03) and the xenogenic silencer Hha was repressed 10-fold (*p*<0.0001) (**Figure 3A-B**). To confirm that HnrS is a direct repressor of Hha we introduced compensatory mutations into the seed-region of HnrS and *hha* 5’ UTR. Single point mutations in either the sRNA or mRNA seed disrupted repression, whereas the compensatory double mutant restored repression, indicating that regulation was dependant on direct base-pairing between HnrS and *hha* RNAs (**Figure 3C-D**).

**Figure 3.**
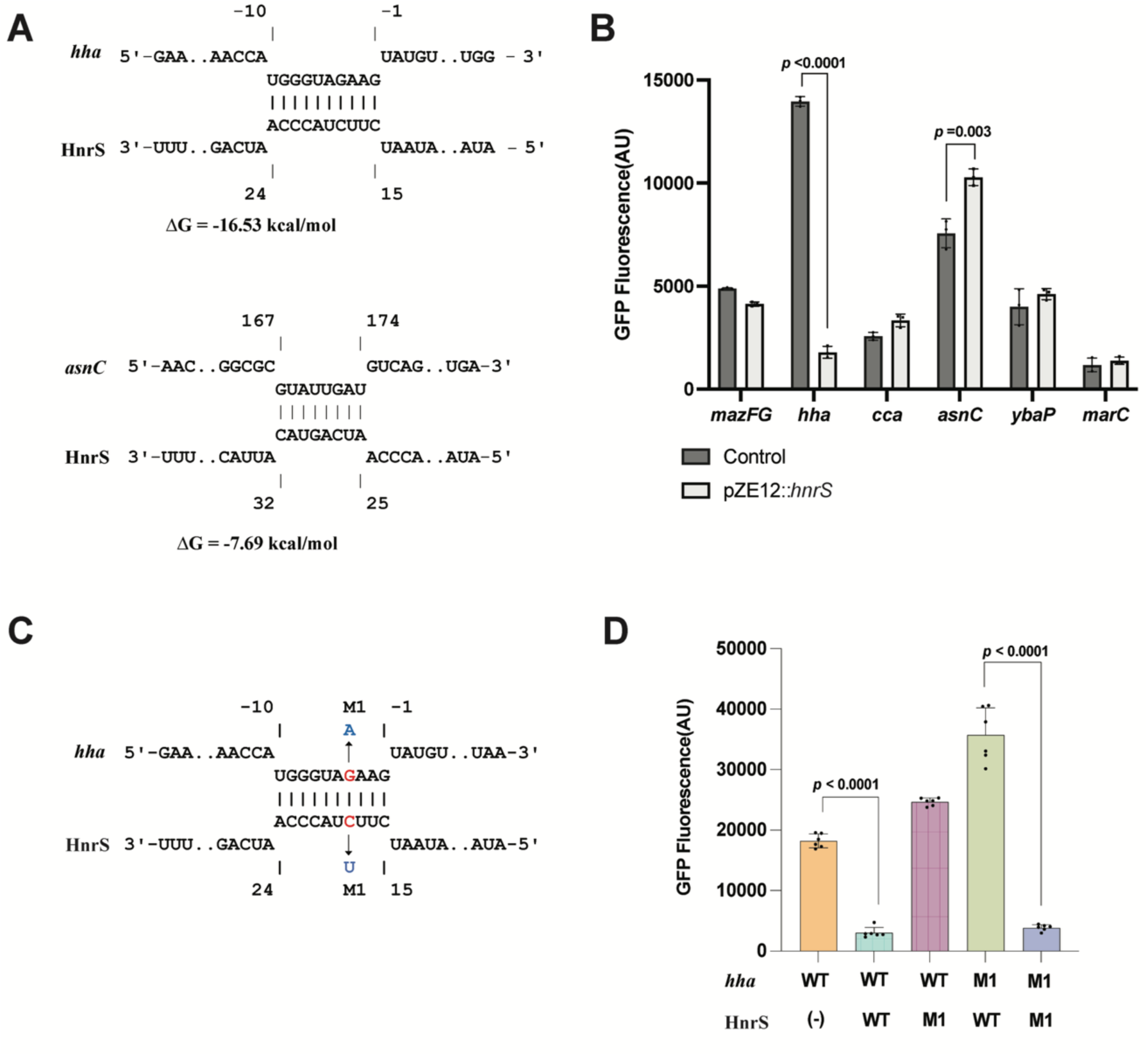
*In silico* and experimental validation of HnrS–mRNA interactions. **A.** Predicted interactions of HnrS with *hha* and *asnC* mRNAs using IntaRNA. **B.** Target mRNA–sfGFP fluorescence in the presence of HnrS or scrambled control plasmid (pJV300) for the first six genes in the RNA interactome. **C.** Compensatory point mutations (blue) were introduced in *hha* or HnrS at the M1 seed region to disrupt predicted base-pairing. The sequence AGAAG upstream of the *hha* coding region corresponds to the ribosome binding site (RBS). **D.** Fluorescence of *hha*–sfGFP was measured in the presence of HnrS wild type or mutant variants, with restoration of interaction by complementary mutations confirming direct pairing. Data represents mean GFP intensity from three biological replicates with two technical replicates each.

Hha forms a complex with H-NS and promotes silencing of predominately horizontally-acquired genes and virulence genes in *E. coli* and *Salmonella* (17, 18, 37). Our data indicate that the prophage-encoded sRNA HnrS represses Hha, suggesting that it acts to counter xenogenic silencing.

### HnrS represses motility in commensal and enterohaemorrhagic *E. coli*

Hha positively regulates motility in *E. coli* and *Salmonella* through activation of the master flagellar regulator FlhDC (38, 39). We initially quantified Hha regulation of motility in EHEC str. Sakai. Single-cell swim speed was measured for the wild type, Δ*hha*, and the complemented mutant (Δ*hha* pBR322::*hha*) using phase-contrast microscopy (**Figure 4A**). Consistent with earlier reports, deletion of *hha* dramatically reduced swim speed in EHEC str. Sakai (4.7-fold, *p*<0.0001) and was restored in the complemented background.

**Figure 4.**
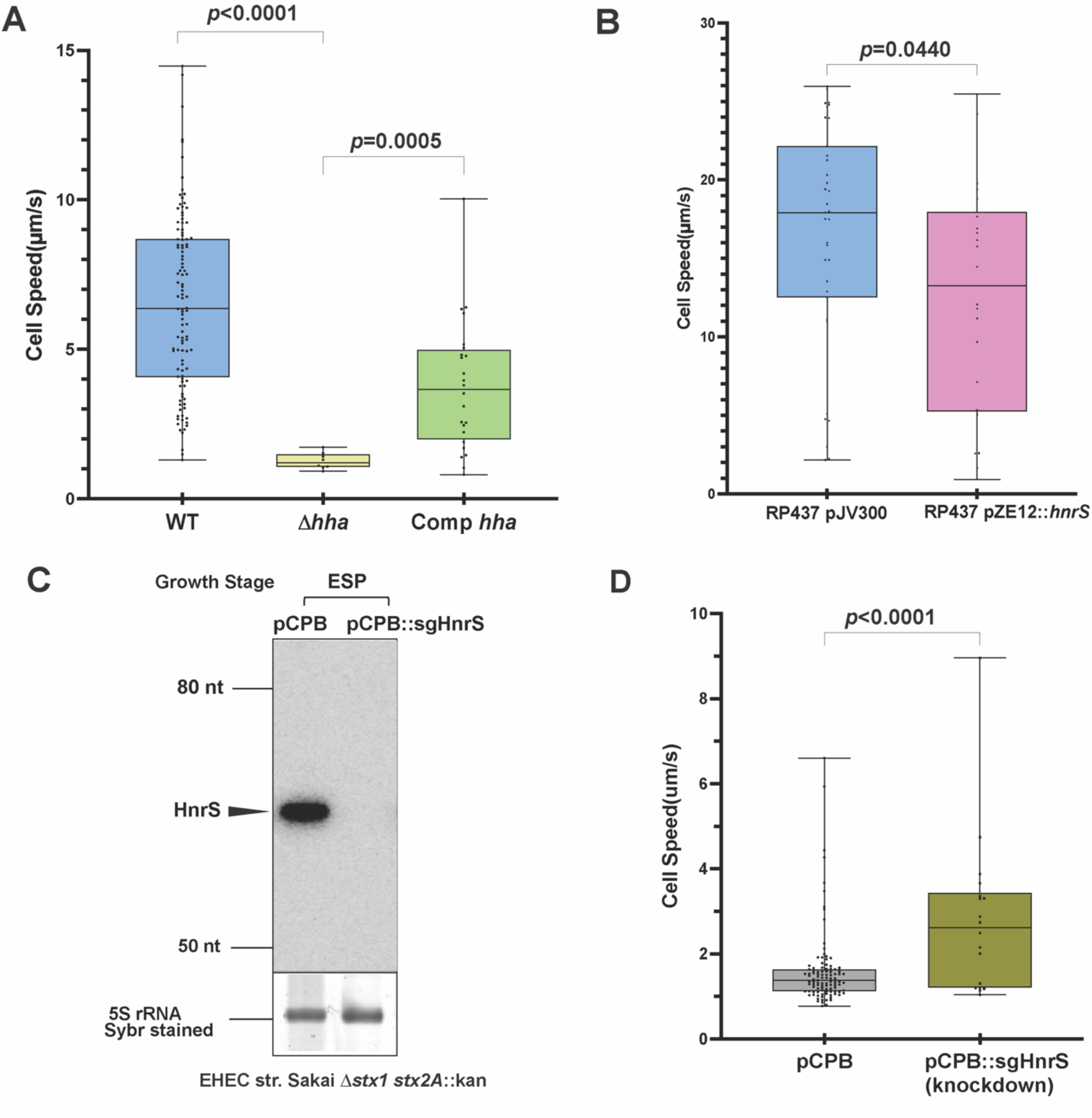
Δ*hha* and HnrS mutants exhibit altered motility. **A.** Single-cell swimming speed of EHEC str. Sakai wild type, Δ*hha* with empty vector (pBR322), and Δ*hha* complemented with pBR322::*hha*. Cells were imaged in tunnel slides under phase-contrast microscopy, and 20-second videos were captured at 20 frames per second (400 frames total) using a Chameleon3 CM3 camera. Speeds <0.5 µm/s were excluded as non-motile. Data were analyzed with LabView and visualised in GraphPad Prism 10 (WT n=118, Δ*hha* n=11, complemented n=30, Mann-Whitney test). **B.** Single-cell speed assay for *E. coli* RP437 wild type, RP437 with empty vector (pJV300), and RP437 carrying pZE12::*hnrS.* **C.** Northern blot detecting HnrS expression in wild type and HnrS CRISPRi knockdown strains (pCPB vector), with RNA harvested at early stationary phase (OD_600_=1.8). **D.** Single-cell speed assay for EHEC Sakai stx(-) strains with empty vector (pCPB) and EcOnc10 knockdown (pCPB::sgHnrS). Videos were captured and analysed as described in panel A.

To assess whether HnrS is able to repress motility, a HnrS expression construct (pZE12::*hnrS*) was moved into the hypermotile commensal *E. coli* str. RP437 (that lacks the *hnrS-*encoding prophages). Expression of HnrS in the commensal background significantly reduced motility compared to a scrambled sRNA control (**Figure 4B**), consistent with HnrS repression of *hha*.

To simultaneously knockdown all three HnrS copies in EHEC we used CRISPR interference (CRISPRi). HnrS was not detectable after knockdown by Northern blot at early stationary phase indicating efficient knockdown (**Figure 4C**). CRISPRi knockdown of HnrS significantly increased motility (1.9-fold, *p=*0.0023), mirroring the overexpression phenotype in commensal *E. coli* str. RP437 (**Figure 4D**).

Because the Δ*hha* mutant is essentially non-motile, epistasis could not be assessed by double mutant analysis. However, our results are consistent with HnrS repressing motility through Hha that is required to activate the FlhDC regulatory cascade.

### HnrS promotes T3SS in EHEC

Expression of the T3SS is essential for EHEC colonisation of both ruminants and humans and is silenced by Hha-H-NS (40–43). Given that the great majority (98.01%) of *hnrS-*encoding *E. coli* are also LEE+, we speculated that a major selective pressure for maintaining multiple copies of HnrS may be de-repression of the LEE to promote colonisation. We initially confirmed that Hha represses the T3SS in our low-secretion EHEC str. Sakai background and in a high-secretion strain, TUV93-0 (a derivative of EDL933 lacking both *stx*-encoding phages). Deletion of *hha* in both strain backgrounds resulted in strong de-repression of T3SS that was restored to wild type levels in the complemented strain (**Figure 5A**).

**Figure 5.**
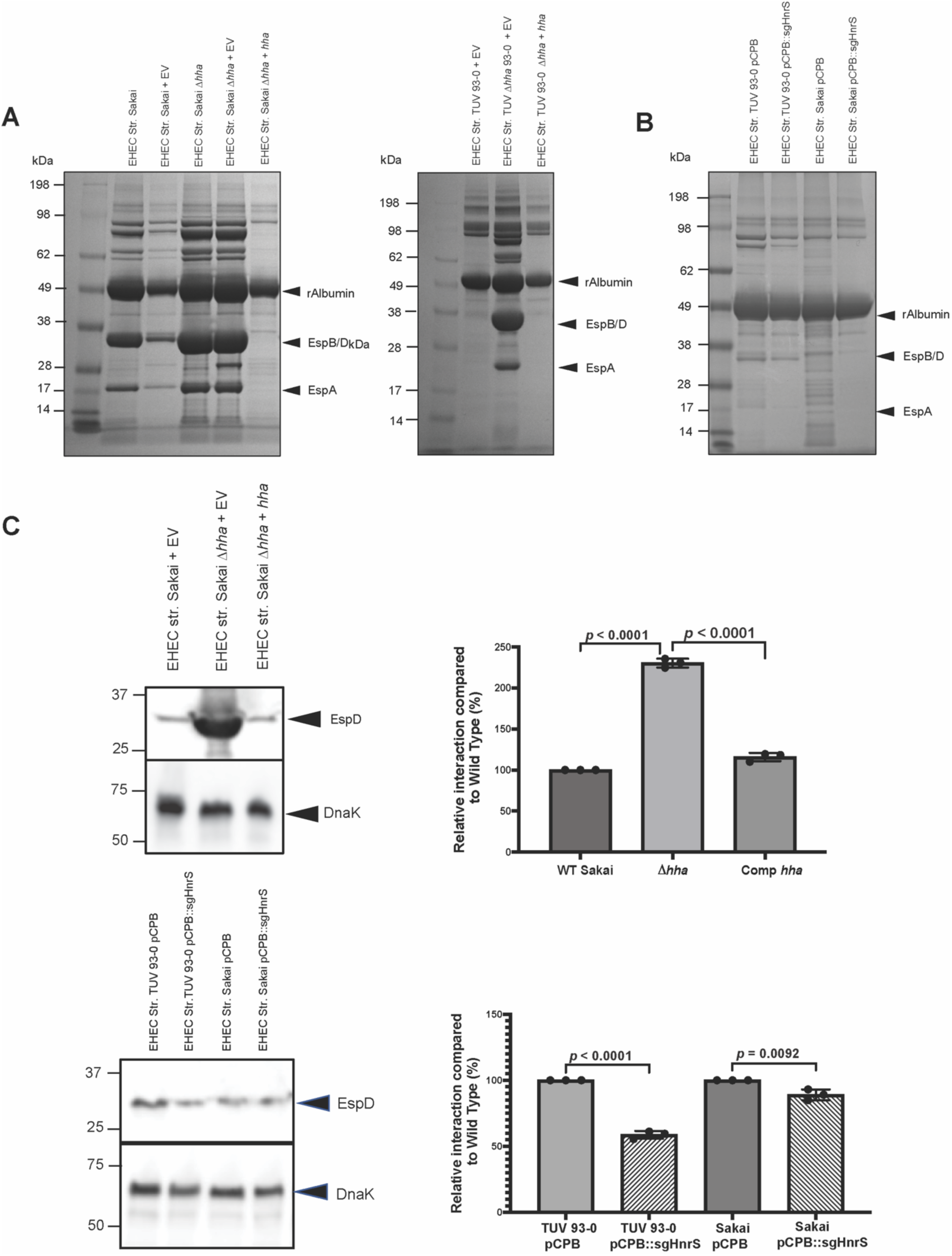
T3SS secretion profiles of hha and HnrS mutants in EHEC Sakai and TUV 93-0 strains. **A.** Coomassie-stained SDS-PAGE gels showing secreted protein profiles. ***Left panel:*** EHEC Sakai stx(-) wild type, wild type with pBR322, Δ*hha*, Δ*hha* with pBR322, and Δ*hha* complemented with pBR322::*hha*. ***Middle panel:*** EHEC TUV 93-0 stx(-) wild type, Δ*hha*, and Δ*hha* complemented with pBR322::*hha.* **B.** Coomassie-stained secreted protein profiles for HnrS knockdown strains. EHEC TUV 93-0 and Sakai strains carrying either the empty vector (pCPB) or pCPB::sgHnrS. Arrows indicate BSA (loading control, 66.4 kDa), EspB/D (needle tip proteins, 32.6 kDa and 39.1 kDa), and EspA (filament, 20.6 kDa). **C.** Western blot analysis of EspD in secreted fractions and DnaK in whole-cell samples. *Top panel:* Sakai derivatives, including wild type with pBR322 (lane 1), Δ*hha* with pBR322 (lane 2), and Δ*hha* complemented with pBR322::*hha* (lane 3). ***Bottom panel:*** Additional strains, including TUV 93-0 with pCPB (lane 1), TUV 93-0 with pCPB::sgHnrS (lane 2), Sakai with pCPB (lane 3), and Sakai with pCPB::sgHnrS (lane 4). Densitometry analysis of the EspD bands from these blots is shown alongside each Western blot. Chemiluminescence images from a technical replicate were quantified using ImageJ. Statistical significance was assessed by unpaired t-test

The T3SS is positively regulated by the master regulator Ler encoded at the start of the LEE1 operon. Using GFP transcriptional fusions to the LEE1 (*ler*), LEE5 (*tir*), and LEE4 (*sepL*) promoters, we quantified promoter activity in the Δ*hha* backgrounds (**Supplementary Figure 4**). Consistent with earlier work (44), deletion of *hha* de-repressed the LEE1 and LEE5 promoters, but not the LEE4 promoter in EHEC str. Sakai and was restored to wild type in the *hha* complementation strain.

CRISPRi knockdown of HnrS reduced the amount of T3S in both strain backgrounds, although baseline secretion levels were lower in the presence of the CRISPRi construct. In Sakai, laddering of cytoplasmic proteins from *stx*-phage lysis obscured identification of the T3SS proteins (**Figure 5B**). To quantify T3SS proteins in the CRISPRi expressing background we used antibodies against the T3SS needle tip, EspD (**Figure 5C**). EspD was de-repressed 2.3-fold, (*p*<0.0001) in the EHEC str. Sakai Δ*hha* background and restored to wild type in the complemented strain. Conversely, *hnrS* knockdown significantly reduced EspD expression 1.7-fold (*p*<0.0001) in the high secretion strain TUV93-0 and 1.1-fold (*p*=0.0092) in the low secretion strain Sakai (**Figure 5D**).

These data confirm Hha repression of the T3SS in EHEC str. Sakai and TUV93-0, and that HnrS increases expression of T3S proteins in these strains, consistent with anti-silencing of the LEE through *hha* repression.

### HnrS increases pedestal formation on epithelial cells

Our results suggest that HnrS expression promotes EHEC towards colonisation through decreased motility and increased T3S-mediated attachment. To assess the impact of HnrS and *hha* on host cell interactions, we incubated the Δ*hha* and HnrS knockdowns strains with embryonic bovine lung (EBLs) epithelial cells, a well-established model for attachment and actin pedestal formation of EHEC (45–48). Despite de-repression of the T3SS that is required for attachment and colonisation, we did not observe a statistically significant difference in total attachment with either the *Δhha* or *hnrS* knockdown strains in the low T3S secretion strain Sakai backgrounds (**Figure 6A**). We next assessed adhesion of the *hnrS* knockdown in the high secretion strain, TUV93-0. As in Sakai, overall attachment to EBL cells was unchanged (**Figure 6B**), but the *hnrS* knockdown had significantly less actin-rich pedestal formation – a key step in T3S-dependant intimate attachment (**Figure 6C**). Quantification at five hours post-infection showed ∼60% of WT TUV93-0 bacteria formed actin-rich pedestals versus ∼30% of (≈1.91 -fold decrease, *p*=0.0079). These data suggest that HnrS increases the rate of intimate attachment in EHEC and is consistent with HnrS priming T3SS expression for attachment and colonisation.

**Figure 6.**
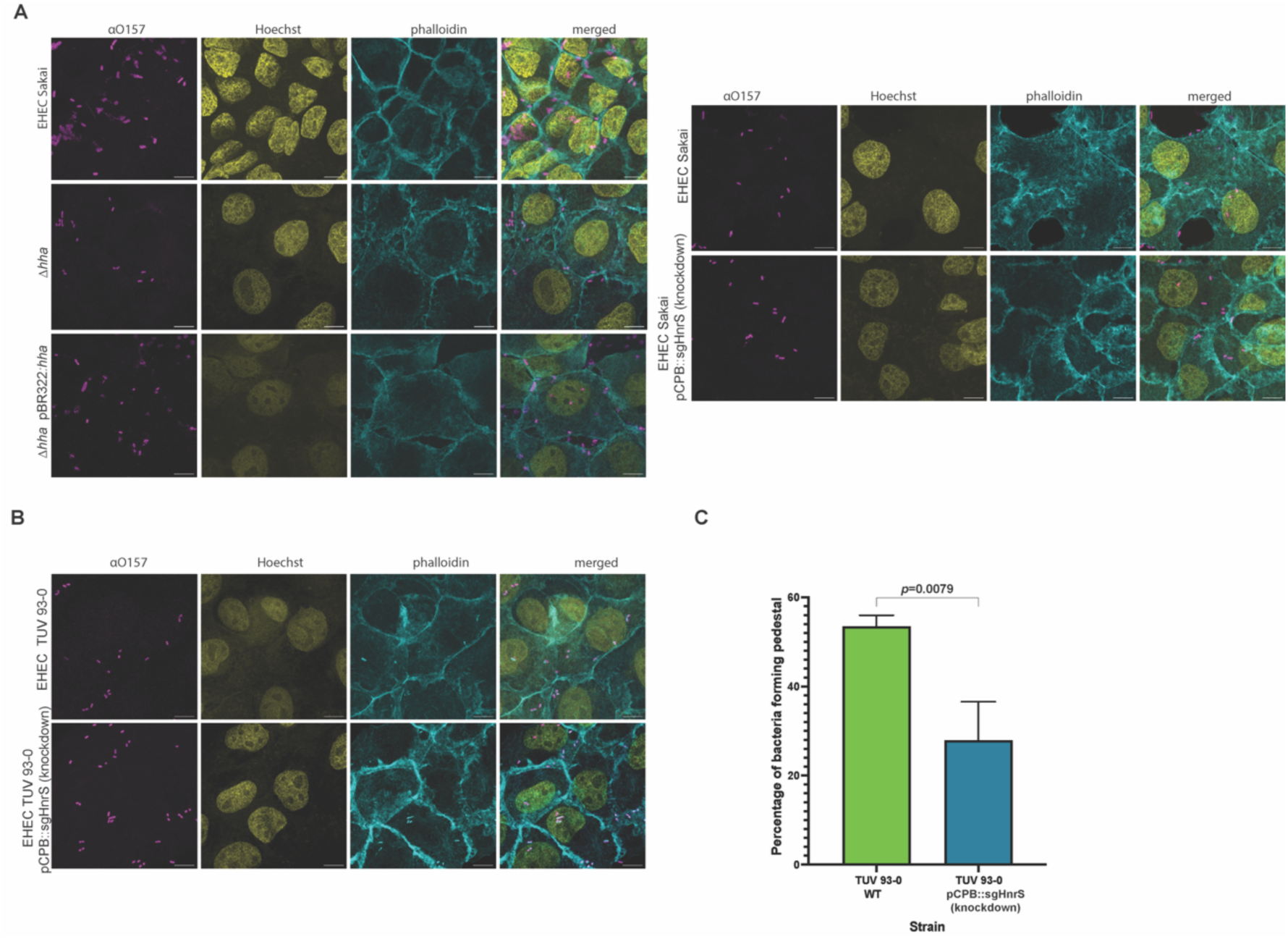
Immunofluorescence analysis of EBL cells infected with EHEC *hha* and HnrS mutants. A. Left panel: EBLs were infected with EHEC Sakai strains carrying pBR322 (wild type), Δ*hha* with pBR322, and the complemented Δ*hha* strain with pBR322::*hha* (MOI 100). Bacteria including flagella (magenta) were visualized using anti-O157 antibodies with Alexa Fluor 568 secondary, nuclei (yellow) were stained with Hoechst 33342, and actin filaments (cyan) were visualized with Phalloidin-FITC. Merged images show co-localization of all stains. Wild type shows intact flagella, Δ*hha* lacks flagella, and the complemented strain restores flagella expression. **Right panel:** EBLs were infected with EHEC Sakai carrying either pCPB (empty vector) or pCPB::sgHnrS (HnrS knockdown) at MOI 100. **B.** EBLs were infected with EHEC TUV 93-0 carrying either pCPB (empty vector) or pCPB:: sgHnrS (HnrS knockdown) at MOI 100. **C.** The percentage of bacteria inducing actin polymerisation at adhesion sites was measured from fluorescence images. TUV 93-0 EcOnc10 knockdown shows significantly reduced actin polymerization compared to the wild type

### HnrS promotes expression of genes involved in anaerobic nitrate/nitrite respiration and nitric oxide resistance

Hha-H-NS silences a broad range of virulence genes in *E. coli* and *Salmonella* (17, 18, 37) suggesting that HnrS may counter-silence other genes required for colonisation. To enable complementation and avoid CRISPRi off-target effects, we constructed a triple knockout,

Δ*hnrS1* Δ*hnrS2* Δ*hnrS3* (hereafter Δ*hnrS*) in the Shiga-toxigenic strain of EHEC str. Sakai using CRISPR–Cas9-mediated genome editing followed by λ-Red recombineering (CRISPathBrick). Deletion of all three copies of *hnrS* was confirmed by Northern blot analysis (**Figure 7A**). The wild-type, Δ*hnrS*, and complemented strains were grown to mid-log phase in T3S-inducing media (supplemented MEM-HEPES). Importantly, HnrS was pulse expressed for 15 minutes before RNA harvesting to primarily capture mRNAs that are direct targets of HnrS. RNA-seq analysis of the transcriptomes indicated that 24-genes were activated in the Δ*hnrS1-3* strain compared to wild type (fold change >2), and 5 were restored to wild type level in the complemented strain **(Supplementary Figure 5)**. While expression levels were low in wild type, *hha* expression was de-repressed in the Δ*hnrS* strain and restored to wild type levels in the complement, consistent with our RNase E-CLASH data and GFP translational fusions (**Figure 7B**).

**Figure 7.**
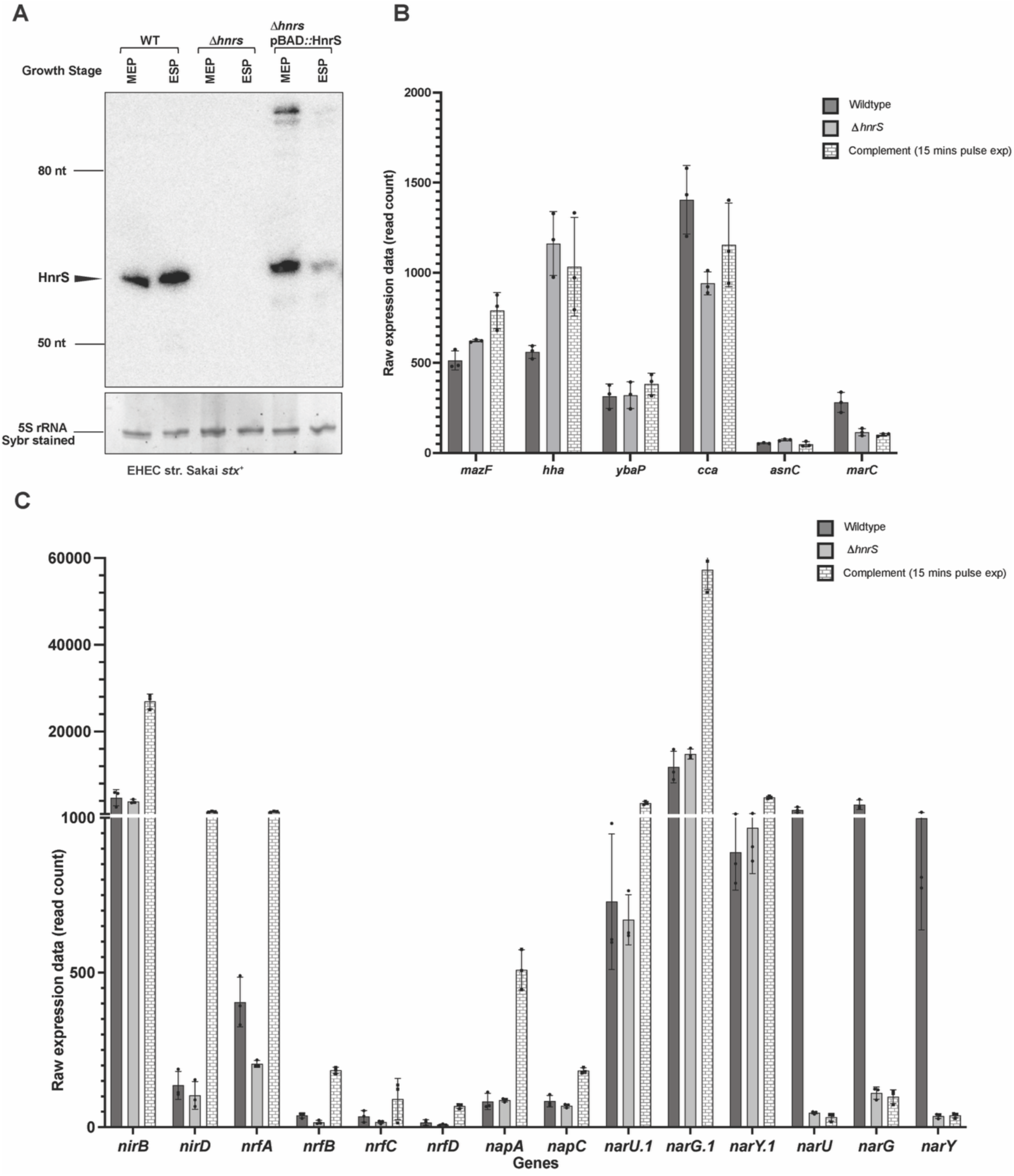
HnrS expression and its impact on nitrate and nitrite respiration genes. **A.** Northern blot showing HnrS expression in wild type (WT), Δ*hnrS*, and complemented strains with pulse expression from pBAD+1::HnrS. RNA was harvested at mid-exponential (OD_600_ = 0.8) and early stationary phase (OD_600_ = 1.8). **B.** Bar graph showing the expression levels of RNase E CLASH targets of HnrS in RNA seq data. Gene expression is quantified by read counts from biological triplicates, based on DESeq data. **C.** Bar graph depicting the expression of genes involved in nitrate and nitrite respiration, showing reduced expression in ΔHnrS relative to WT and complemented strains. Gene expression is quantified from DESeq2 read counts of biological triplicates. *narU.1*, *narG.1*, and *narY.1* correspond to *E. coli* O157:H7 Sakai homologues that are not present in *E. coli* K-12 MG1655, although they share nomenclature with *narU*, *narG*, and *narY*.

Ontological clustering indicated that genes involved in nitrogen metabolism were significantly regulated by HnrS (**Supplementary Figure 6**), including HnrS activation of the *nirBD, nrfABCD*, *napA* and *narGU* operons involved in nitrite and nitrate respiration (**Figure 7C**). The nitric oxide reductase encoded by *hcp* was significantly activated by HnrS in the Δ*hnrS* strain and was the most upregulated after pulse expression of HnrS (**Figure 7C**), indicating that HnrS expression promotes expression of nitrite/nitrate respiration genes and nitric oxide detoxification.

To identify direct targets of HnrS in our differentially expressed genes, we predicted base-pairing complementarity with HnrS and constructed GFP translational fusions for several differentially expressed genes (**Supplementary Figure 7**). Among the nitrogen metabolism genes with complementarity (*nirB*, *nirD*, *hcp*, *nrfA*, *napF*, *narQ*), none were significantly upregulated by HnrS, suggesting that regulation of nitrogen metabolism is indirect.

To understand how much of this regulation can be explained by HnrS regulation of *hha*, we compared the differentially expressed genes to RNA-seq data from EHEC Δ*hha* (43). Hha regulates a large proportion of the EHEC genome and a substantial subset of genes that were differentially regulated by *hnrS* deletion or pulse expression were also regulated by Hha, (**Supplementary Figure 8**), including several genes involved in nitrogen metabolism (*nirBC*, *napGHJI*, *napBC*). These data suggest that HnrS may act indirectly (at least partly) to regulate nitrogen metabolism operons through repression of Hha.

During gastrointestinal colonisation, inflammation drives epithelial responses and nitric oxide is secreted by host macrophages to produce NO, which is rapidly oxidised to nitrite and nitrate; these anions accumulate and function as terminal electron acceptors for anaerobic respiration of *E. coli* (24). By repressing *hha*, HnrS appears to promote expression of many nitrate/nitrite respiration and nitric oxide resistance genes required to prime EHEC for survival in the inflamed gastrointestinal tract, at least partly through counter-silencing of *hha*. Together, the data suggest that HnrS primes EHEC for gastrointestinal colonisation by counter-silencing pedestal formation (T3SS) and anaerobic nitrate/nitrite respiration genes while repressing motility.

## DISCUSSION

H-NS silencing of foreign DNA serves as a final layer of genome defence and allows the cell to balance protecting the genome with acquisition of new genetic information. H-NS silencing is enhanced at horizontally-acquired virulence genes in *E. coli* and *Salmonella* by the H-NS paralogues Hha and YgdT (17, 18, 49). To overcome silencing, many virulence loci encode transcription factors like Ler, Sly, and VirB, that act narrowly to counter-silence specific virulence genes in response to genetic or environmental signals, allowing the virulence genes to be positively selected (15, 50). Complete de-repression of H-NS is likely to have severe fitness costs and deletion of *hns* in *Salmonella* and *Yersina* is lethal (17, 18, 37, 51). H-NS antagonists like the truncated H-NS paralogue, H-NST, are expressed at low levels to avoid full H-NS de-repression (51, 52). Here we demonstrate that the prophage-encoded sRNA, HnrS, relieves Hha-H-NS silencing of the T3SS in EHEC through direct post-transcriptional repression of Hha by base-pairing at its ribosome binding site. We propose that counter-silencing through Hha, rather than H-NS, allows more specific de-repression of horizontally-acquired virulence genes and avoids fitness costs associated with de-repression of the core H-NS regulon.

We find that HnrS counter-silencing allows EHEC to form more actin-rich pedestals on cultured epithelial cells suggesting that it primes EHEC for gastrointestinal colonisation. This is consistent with down regulation of motility, upregulation of T3SS, and upregulation of genes required for anaerobic nitrate/nitrite respiration and nitric oxide resistance. During mucosal inflammation, epithelial cells and recruited myeloid cells (macrophages and neutrophils) induce inducible nitric oxide synthase (iNOS), generating NO that is rapidly oxidised into nitrates and nitrites (24, 25). These anions accumulate at the hypoxic mucosal surface and function as terminal electron acceptors for anaerobic respiration by *E. coli*. These electrons receptors are used for anaerobic respiration as demonstrated by attenuation of a *napG* mutant encoding nitrate reductase (24, 25). Deletion of *hnrS* reduced expression of several pathways for nitrate and nitrite respiration including *napG* and the NO-detox system *hcp/hcr*. In EHEC, nitrates also trigger maturation of the T3SS by acting as a checkpoint between the basal body and needle filament assembly (24) providing an additional layer of coordination between T3SS and gastrointestinal colonisation. Because HnrS does not directly regulated reporters for several nitrogen metabolism genes, their regulation is likely indirect and *hha*-dependant, although we cannot rule out direct regulation through an untested pathway. Interestingly, *hcp* is strongly upregulated after 15 minutes of HnrS transcription suggesting direct regulation. Collectively, our data indicate that HnrS primes a programme required for gastrointestinal colonisation through counter-silencing of Hha, shifting from a motile/dispersal state towards an attached, pedestal forming, and nitrate-respiring state.

The HnrS sRNA is encoded on prophages and this has led to repeated acquisition of the sRNA. Both EHEC strains Sakai and EDL933 carry three copies on separate prophage. In some EHEC and EPEC strains *hnrS* has been acquired up to nine times. Additional copies of *hnrS* likely increase expression through gene dosage and we predict that those strains with high copy numbers are likely to have higher levels of counter-silencing. Interestingly, despite being encoded on a mobile genetic element, *hnrS* is almost exclusively found in LEE+ *E. coli* strains, further supporting a function for HnrS in attaching and effacing lesion formation.

While we have demonstrated that HnrS modulates EHEC virulence gene expression it is less clear whether HnrS provides a selective advantage to the phage. HnrS interacts with the toxin-antitoxin (TA) system *mazEF* in our *in vivo* sRNA interactome data and *in vitro* by EMSA, but we were unable to define a phenotype for this interaction. Several TA systems confer phage resistance through an abortive infection mechanism (53) and MazEF is reported to confer resistance to phages P1 and T4 (34, 35), but we were unable to reproduce this phenotype, or demonstrate HnrS regulation of MazF toxicity or *mazG* expression. A recent study has indicated that MazEF confers resistance to RNA phages (54). Unexpected new functions for sRNAs in regulating co-translational protein folding through interactions with the mRNA coding sequence have recently been uncovered (36). Given the unusual position of the HnrS-*mazF* interaction (perfect complementary to the last 15-nt of the *mazF* coding sequence), we speculate that HnrS may be functioning by a non-canonical mechanism that is not captured by our assays.

Prophages form one of major building blocks of pathogen evolution and overcoming xenogenic silencing is required for expression and positive selection of virulence genes encoded by the phage. Several mobile elements carry truncated, dominant negative paralogues of H-NS to relieve silencing (16, 52). Here, we find that prophage utilise a regulatory sRNA that counters Hha silencing. HnrS confers Hha-specific counter silencing that likely avoids disrupting the core H-NS regulon. Regulatory sRNAs are compact and highly portable. We predict other mobile genetic elements will encode sRNAs that relieve xenogenic silencing to positively select their virulence gene cargo.

## METHODS

### Bacterial strains and growth conditions

Bacterial strains, primers, plasmids and probes used for this study are listed in Supplementary Table 1. *E. coli* was routinely cultured at either 30°C or 37°C in LB broth, minimal M9 broth, MEM-HEPES (Sigma M7278) supplemented with 0.1% glucose and 250 nM Fe(NO_3_)_3_, or solid LB agar plates. Bacterial media was supplemented with ampicillin (100 μg/mL), chloramphenicol (34 μg/mL), kanamycin (50 μg/mL), tetracycline (10 μg/mL) or spectinomycin (50 μg/mL) where appropriate.

### Construction of EHEC deletion strains

Chromosomal deletions of *hha* and *mazEFG* were constructed using the pTOF25 allelic exchange system (55). Deletion fragments were generated by SOE-PCR and cloned into pTOF25 carrying an FRT–tetRA–FRT cassette for selection. Recombinants were isolated following allelic exchange, confirmed by PCR and sequencing, and the resistance cassette was excised using FLP recombinase (Supplementary Methods).

### CRISPRi knockdown of sRNA HnrS

The pCRISPathBrick (pCPB) vector (Addgene, USA) was used for targeted knockdown of *hnrS* in *E. coli* Sakai. sgRNAs targeting *hnrS* were designed, phosphorylated, and annealed. The pCPB vector and sgRNA inserts were digested with BsaI and ligated to generate pCPB::HnrS. The verified plasmid was electroporated into *E. coli* Sakai stx(-) and selected on LB agar with kanamycin and chloramphenicol.

### Construction of HnrS deletion in EHEC str. Sakai using CRISPR-Cas9

The HnrS gene in *E. coli* O157:H7 strain Sakai stx⁺ was deleted using a CRISPR-Cas9-based system, following a two-plasmid approach (56). The *hnrS* gene in *E. coli* O157:H7 Sakai stx⁺ was deleted using a CRISPR-Cas9 system with a two-plasmid approach (57). A single guide RNA (sgRNA) targeting *hnrS* was cloned into pTarget-F. Homology arms (∼300 bp upstream and downstream of *hnrS*) were joined via SOE PCR and inserted into the pTarget-F vector to generate pTargetT::*hnrS*. Plasmids were transformed into DH5α for propagation and sequence verification. For genome editing, pCas expressing the λ-Red recombination system was introduced into Sakai stx⁺, induced with L-arabinose at mid-exponential phase, and pTargetT::*hnrS* was electroporated. Transformants were selected on LB agar with kanamycin and spectinomycin, and deletion of *hnrS* was confirmed by colony PCR. Positive colonies were cured of the editing plasmid by growth on LB-kanamycin with IPTG at 42°C.

### Phage plaquing assay

Plaque formation and phage titres were determined using the double-agar overlay method (58, 59). LB bottom agar (2% agar) was poured into 10 × 10 cm plates and allowed to dry. Top agar (0.5% agar) was supplemented with 10 mM CaCl₂, 10 mM MgSO₄, and 800 µL of overnight *E. coli* MG1655 cultures (with or without EcOnc10); for Stx phage assays, 1.5 µg/mL mitomycin C was included. Serial dilutions of phages were spot-plated on the top agar, and plates were incubated overnight at 37°C. Plaques were counted the following day to determine phage titres.

### Confirmation of sRNA-mRNA interactions using the sfGFP 2-plasmid system

Prediction of RNA-RNA interaction was done using computational tool IntaRNA (60). GFP translational fusions were generated following established protocols (61, 62). Detailed cloning procedures, including primer sequences and amplification conditions, are provided in Supplementary Methods. In brief, pXG30::*insert* or pXG10::*insert* were co-transformed with either pZE12::*hnrS* or pJV300 (pZE12 encoding a scrambled RNA sequence) into E. coli DH5α. Biological triplicates were grown overnight in LB supplemented with chloramphenicol and ampicillin, overnight cultures were diluted into 15 mL of filtered LB broth supplemented with chloramphenicol and ampicillin, and incubated until OD₆₀₀ reached 0.6. The GFP fluorescence of GFP fusion plasmids was assessed using a POLARstar Omega microplate reader. Fluorescence measurements were taken at an emission wavelength of 510 nm, with an excitation wavelength of 485 nm, which is optimal for GFP fluorescence.

### Northern Blot

Total RNA was extracted from *E. coli* O157:H7 str. Sakai using a guanidinium thiocyanate– phenol method (63) (see Supplementary Methods for details). Five micrograms of RNA were separated on 8% denaturing polyacrylamide gels and transferred to nylon membranes. RNA was immobilised by UV-crosslinking, and specific sRNAs were detected using ^32^P-labelled oligonucleotide probes. Hybridisation and washes were performed under standard conditions, and signals were visualised using a phosphorimager (Typhoon™ FLA9500). Detailed protocols are provided in the Supplementary Methods.

### RNA sequencing analysis for gene expression

*E. coli* str. Sakai and isogenic strains grown in MEM-HEPES (Sigma M7278) supplemented with 0.1% glucose and 250 nM Fe(NO_3_)_3_ at 37°C with shaking until OD_600_ 0.8 and RNA extraction was performed as above. Following RNA extraction, samples were treated with RQ1 RNase-Free DNase (Promega, Cat. no.: M6101) in the presence of RNasin Ribonuclease inhibitor (Promega, Cat. no.: N2511) for 30 minutes at 37 °C to remove residual genomic DNA. RNA was then purified and concentrated using the Zymo RNA Clean & Concentrator kit (Zymo, Cat. no.: R1019), and quality and concentration were assessed using a Nanodrop spectrophotometer and an Agilent TapeStation 4150 to determine RINe values.

The prepared RNA samples were sent to NovogeneAIT Genomics, Singapore for paired-end 150 bp sequencing using the NovaSeq platform. Sequencing reads were processed with the READemption v2.0.4 pipeline, aligning reads to the *E. coli* O157:H7 Sakai reference genome (accession number NC_002695.2) and quantifying transcript abundance. Differential gene expression analysis was carried out with a minimum Phred quality score of 30 to ensure high-confidence data. RNA sequencing datasets are available at NCBI GEO under the accession GSE311113.

### HnrS prevalence analysis in *E. coli* genomes

Genomic data for 3,230 *E. coli* strains were retrieved from NCBI and analysed to assess strain identity, serotype, and pathovar classification. Key housekeeping genes (*recA*, *purA*, *mdh*, *icd*, *gyrB*, *fumC*, *adk*) and virulence/pathotyping markers were identified using K-mer Analysis (KMA) and Shigella/ ST(E)C Finder tools. Boolean logic rules were applied to classify strains into pathovars such as EHEC, EPEC, EAEC, UPEC, and others. The relative coverage of the HnrS was calculated from KMA output by normalising its depth to housekeeping genes, providing an estimate of gene copy number across strains. Detailed methods are provided in Supplementary Methods.

### Determination of bacterial motility

Bacterial strains were grown on LB-agar plates and single colonies were subcultured in LB broth to mid-exponential phase. Motility was initially assessed on low-agar plates (0.2% agar) by measuring the diameter of bacterial flares. For single-cell analyses, cultures were diluted to approximately 1 × 10⁵ cells/mL and introduced into flow cells for imaging (64). Bacterial movement was recorded using phase-contrast microscopy with a 40× objective, capturing 20-second time-lapse videos at 20 frames per second. Swimming speeds were determined using custom LabVIEW software(65) and the resulting data were plotted and analysed using GraphPad Prism 8.

### Type III secretion assay and immunoblotting

Overnight LB cultures were diluted 1:100 into MEM–HEPES with 250 nM Fe(NO₃)₃ and 0.1% glucose and grown to OD₆₀₀ ≈ 0.8. Cultures were centrifuged, and supernatants were filtered (0.45 μm) and precipitated overnight at 4°C with 10% TCA, using recombinant albumin as a co-precipitant (66). Pellets were resuspended in SDS-PAGE buffer, heated, and separated on 4–20% gradient gels, which were stained with Coomassie and imaged.

For immunoblotting, proteins were transferred to nitrocellulose, blocked with 8% milk in PBS, and EspD was detected using mouse anti-EspD and HRP-conjugated secondary antibodies with ECL. DnaK in whole-cell lysates was used as a loading control under the same conditions.

### Cell adhesion assay

Bovine embryonic lung epithelial cells (EBLs) were seeded at ∼10⁵ cells/well in 24-well plates and incubated overnight at 37°C with 5% CO₂. Bacterial strains grown overnight in LB were subcultured 1:10 in MEM–HEPES with 0.1% glucose and 250 nM FeNO₃ for ∼3 h. One hour before infection, EBL media was replaced with MEM–HEPES, and cells were infected at an MOI of ∼100. At 3 and 5 h post-infection, wells were washed, and adhered bacteria were recovered by trypsin and quantified by serial dilution plating on LB agar.

For microscopy, collagen-coated coverslips were fixed with 1% paraformaldehyde for 10 min, permeabilised with 0.1% Triton X-100 for 10 min and blocked with 1 mg/mL bovine serum albumin for 1 h at room temperature. Cells were incubated with rabbit anti-O157 primary antibody (1:50) for 1 h, followed by Alexa Fluor 568–conjugated goat anti-rabbit IgG secondary antibody (1:1000) for 1 h in the dark. Actin filaments were labelled with Phalloidin– Alexa Fluor 488 (BioLegend) for 30 min, and nuclei were counterstained with Hoechst 33342 (1:2000) for 5–10 min. Coverslips were mounted in ProLong Antifade and imaged on a Zeiss LSM900 confocal microscope (63× oil objective) using 488, 568, and 405 nm excitation. Detailed staining procedures are provided in the Supplementary Methods.

### Data availability

RNA sequencing datasets generated in this study are available at NCBI GEO under the accession GSE311113.

## SUPPLEMENTARY FIGURES

**Supplementary Figure 1.**
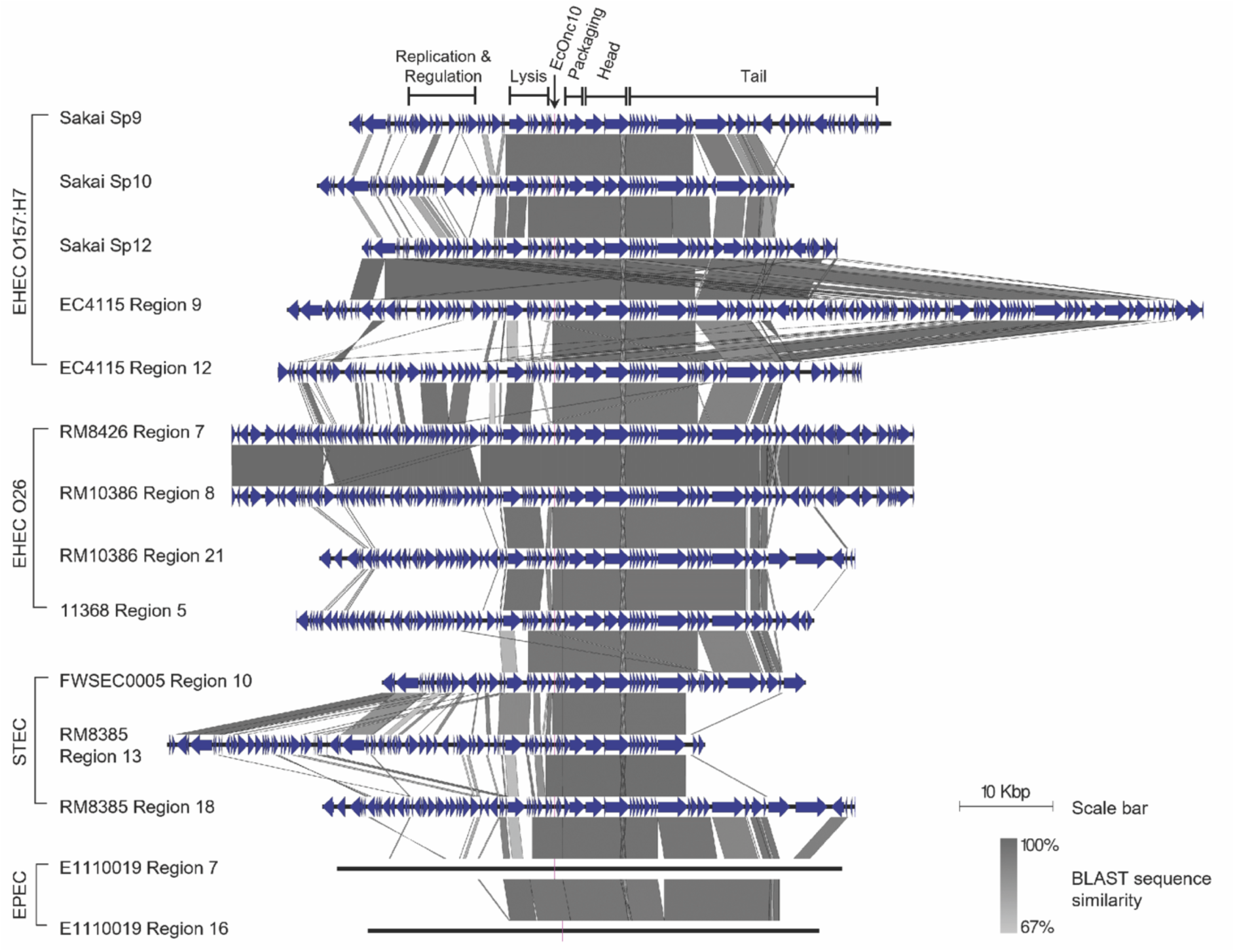
HnrS is encoded within prophages in E. coli pathotypes. Prophage regions from eight different E. coli pathotypes were aligned based on sequence similarities between their prophage genomes as determined by BLAST. HnrS is indicated in pink and labelled as EcOnc10 in the alignment. Annotations for phage genes and hypothetical proteins are shown in blue and were retrieved from NCBI; however, no annotations are available for the prophage regions of the EPEC strain E1110019. On the left, the pathotype, serotype, and strain identifiers are listed, followed by either the name of the prophage (for strain Sakai) or the region number assigned by PHASTER. For strain Sakai’s prophage Sp9, the regions responsible for phage replication, regulation, lysis, packaging, head, and tail fibre genes are marked.

**Supplementary Figure 2:**
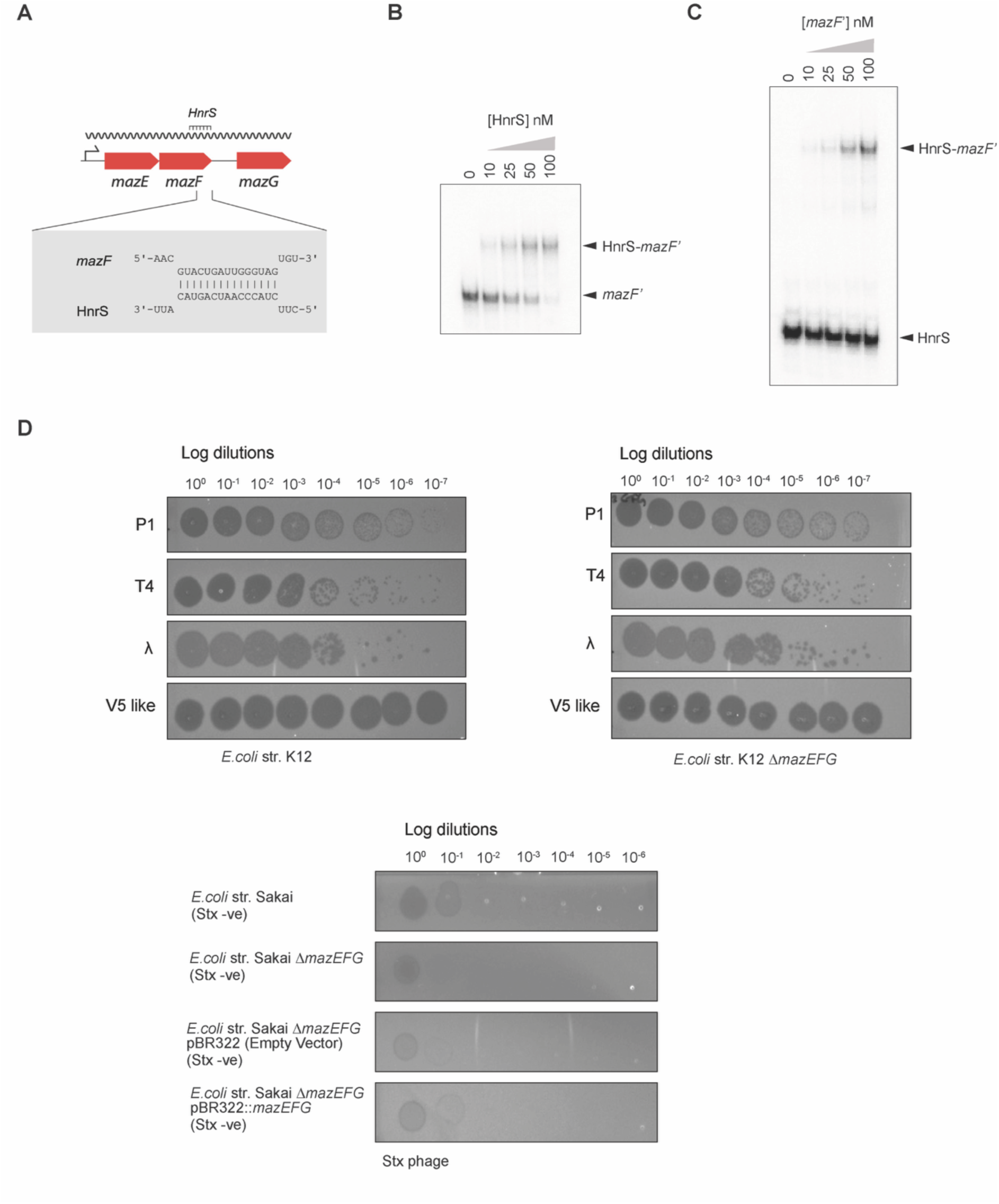
HnrS interaction with the MazEF toxin-antitoxin system and phage sensitivity assays. **A.** *In silico* prediction of HnrS binding to the *mazF* mRNA, showing the predicted interaction site along with neighbouring genes in the *mazEFG* operon. **B.** EMSA to show HnrS interacts with *mazF* coding sequence invitro. **C.** Top. Plaquing assay to assess the sensitivity of *E. coli* K12 wild type (top) and Δ*mazEFG* (bottom) to a panel of phages, including P1kc, T4-like, λ, and V5. Serial dilutions of each phage were spot plated onto the respective bacterial strains, and phage plaques were counted to assess bacterial sensitivity. Bottom: Sensitivity of EHEC strain Sakai, Sakai Δ*mazEFG*, and complemented strains (overexpressing *mazEFG* from pBR322) to Stx phage induction (bottom).

**Supplementary Figure 3:**
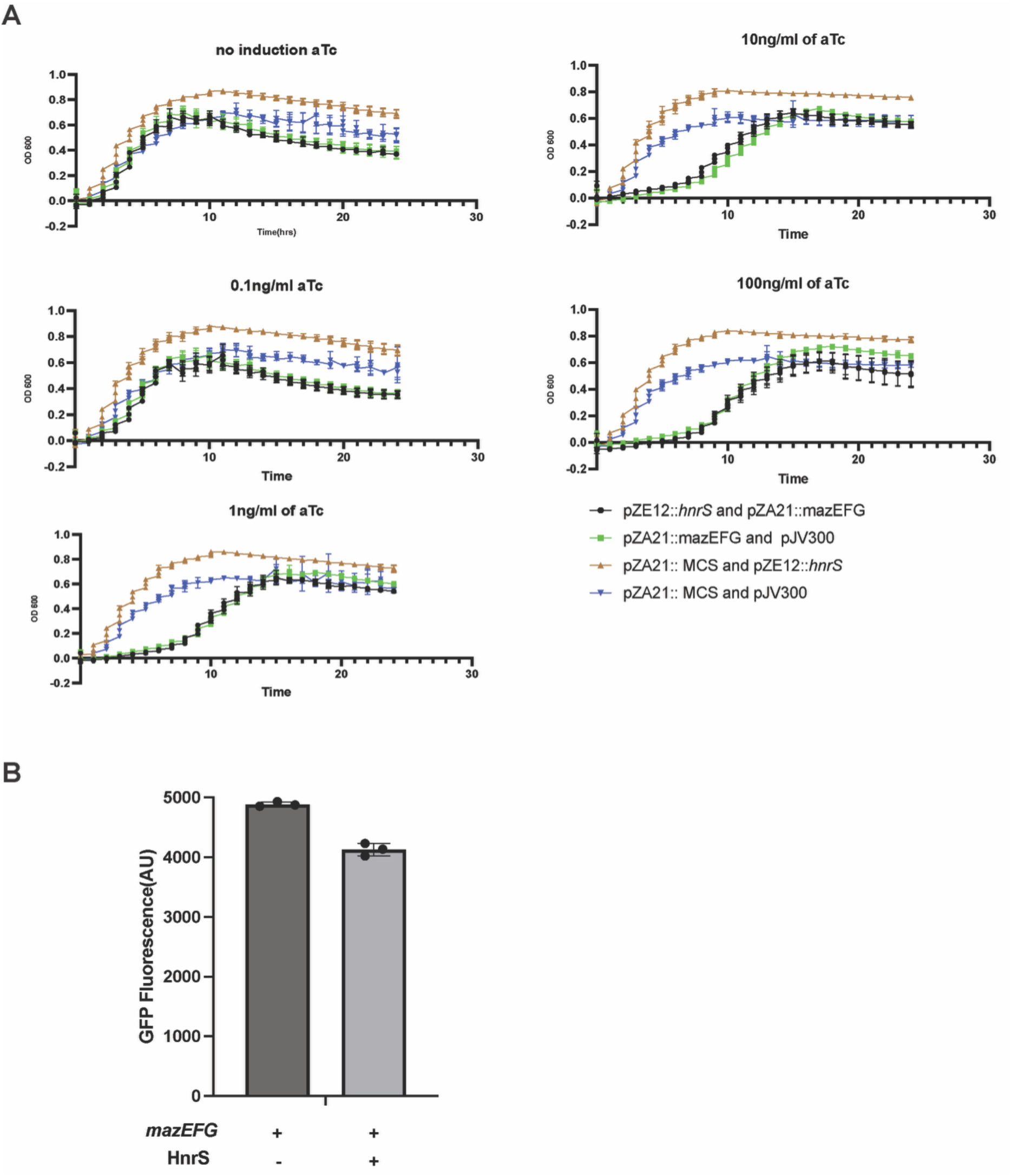
HnrS does not repress MazEF toxicity in *E. coli.* **A.** Growth curves of *E. coli* Top10F’ overexpressing HnrS (pZE12::HnrS) and/or MazEFG (pZA21::mazEFG) under different induction conditions. Cultures were grown at 37°C with shaking for 24 hours in a Bioscreen C system. Panels show growth without aTc (no MazEFG induction) and with increasing aTc concentrations (0.1, 1, 10, 100 ng/mL) to induce MazEFG expression. Control plasmid combinations are indicated as described in Materials and Methods. **B.** Fluorescence of a MazEFG-sfGFP translational fusion in *E. coli* DH5α under different expression conditions of HnrS. Error bars represent standard deviations from biological replicates.

**Supplementary Figure 4.**
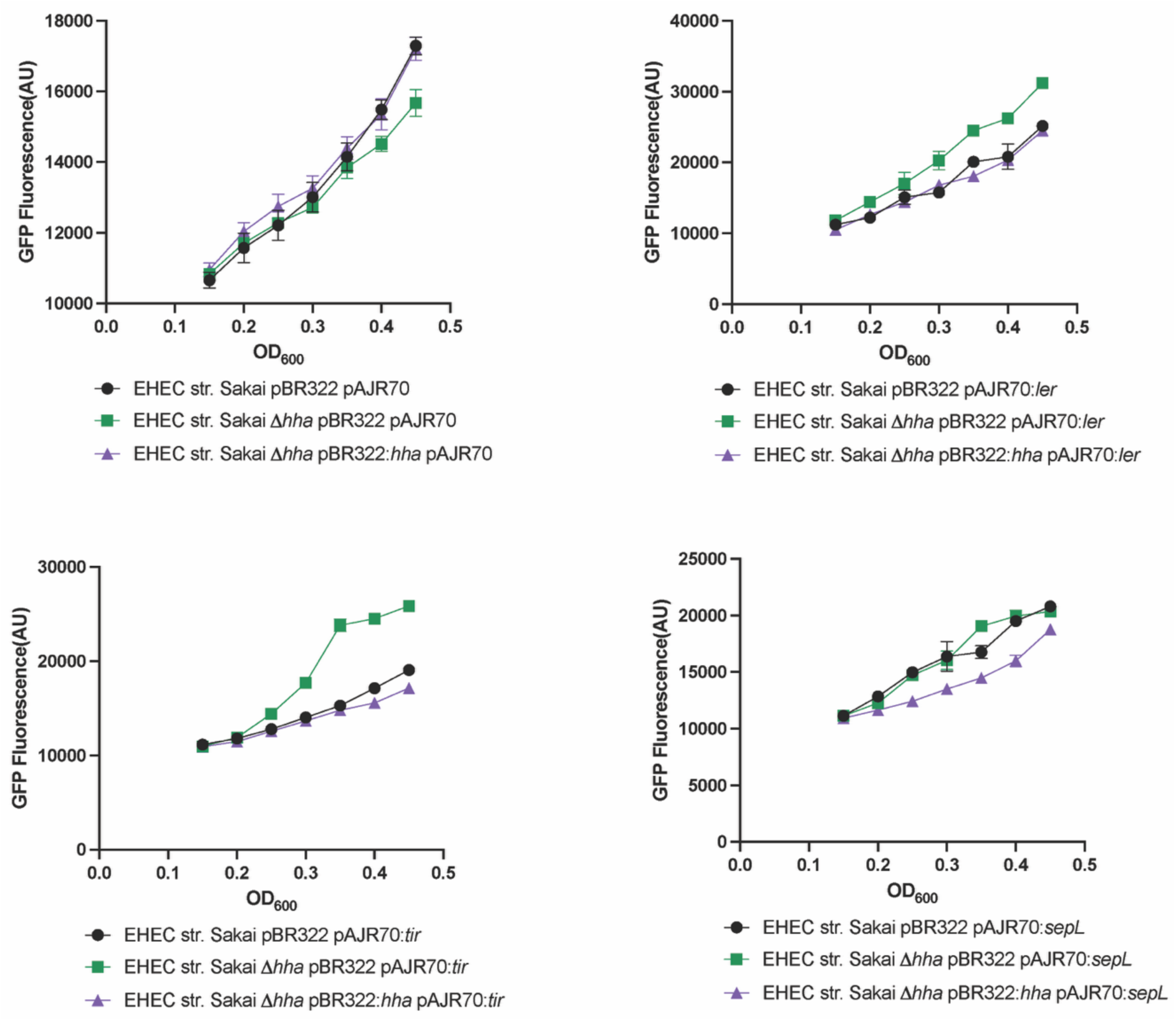
GFP fluorescence versus OD600 for LEE promoter fusions. Individual LEE operon promoters (*ler*, *tir*, *sepL)* were fused to GFP and introduced into EHEC Sakai strains carrying either pBR322 (wild type), Δhha with pBR322, or Δhha complemented with pBR322::hha. The promoter-less GFP plasmid pAJR70 served as a control. Fluorescence (GFP) and OD600 were recorded every 20 minutes. Data points represent the mean of three biological replicates, each with two technical duplicates; error bars indicate ±1 SD. A representative experiment of at least three independent trials is shown.

**Supplementary Figure 5.**
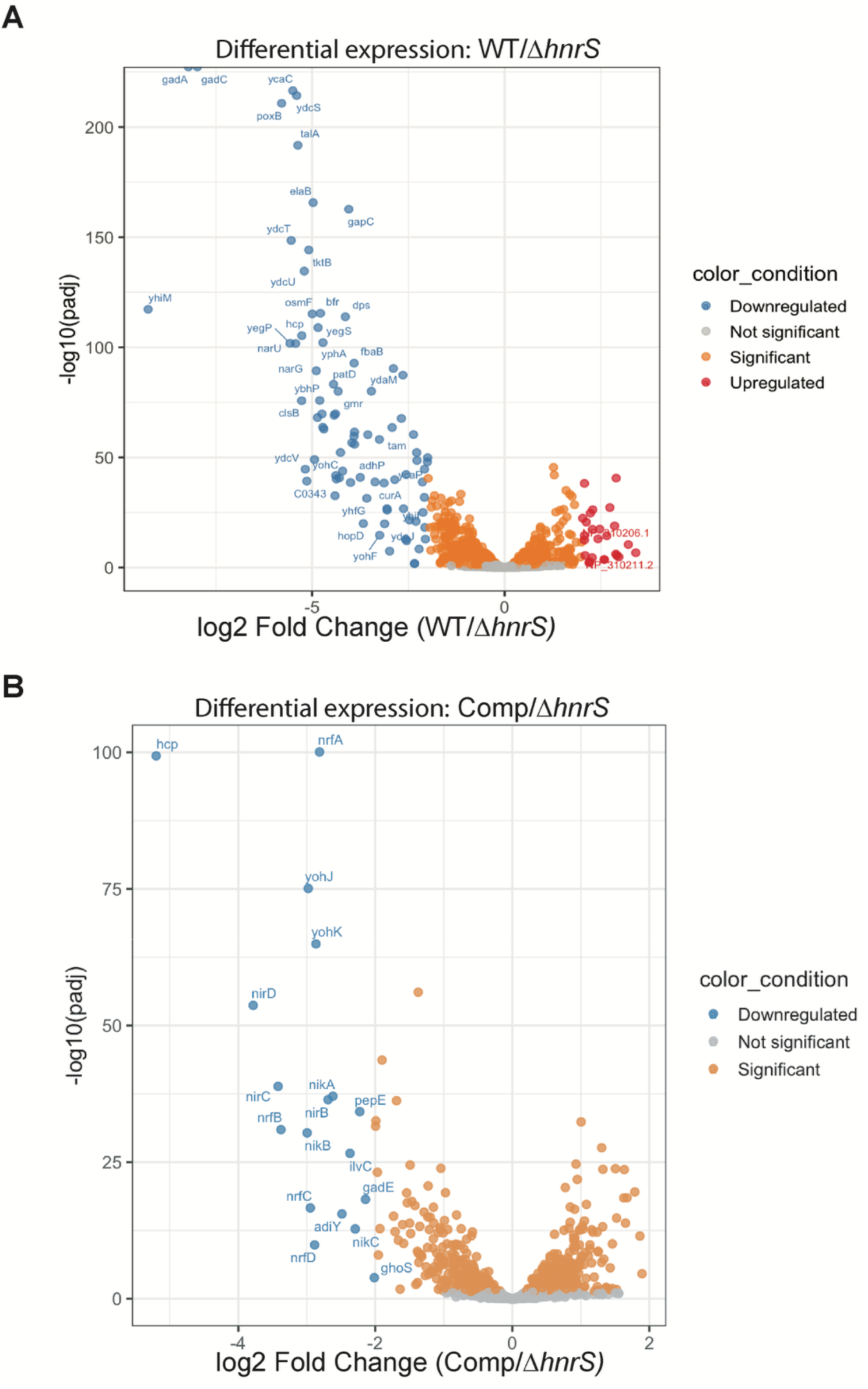
Differential gene expression analysis of EHEC Sakai strains under wild type, deletion, and complementation of HnrS. A: Volcano plot showing differentially expressed genes between the deletion mutant (EHEC Sakai ΔHnrS) and the wild type (fold change ≥ 2, p < 0.05). **B:** Volcano plot comparing the deletion mutant (EHEC Sakai ΔHnrS) to the complementation strain (EHEC Sakai ΔHnrS with pBAD+1::*hnrS*). Green and red dots indicate significantly upregulated and downregulated genes, respectively.

**Supplementary Figure 6.**
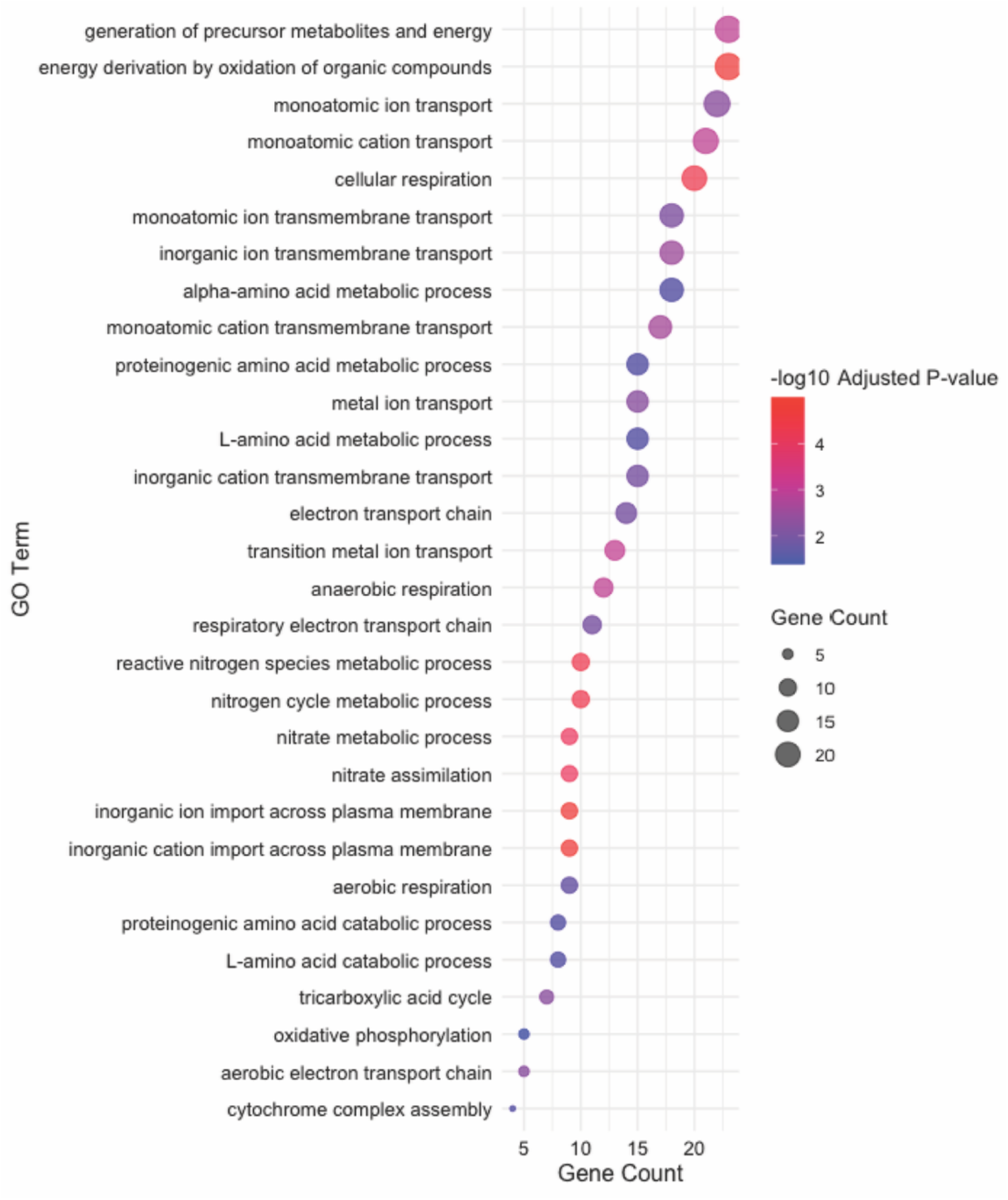
Gene Ontology (GO) analysis of differentially expressed genes (DEGs) between the ΔHnrS mutant and wild type strain. GO analysis under the Biological Process category identified processes significantly enriched (p-adjusted < 0.05) in the wild type strain compared to the ΔHnrS mutant.

**Supplementary Figure 7.**
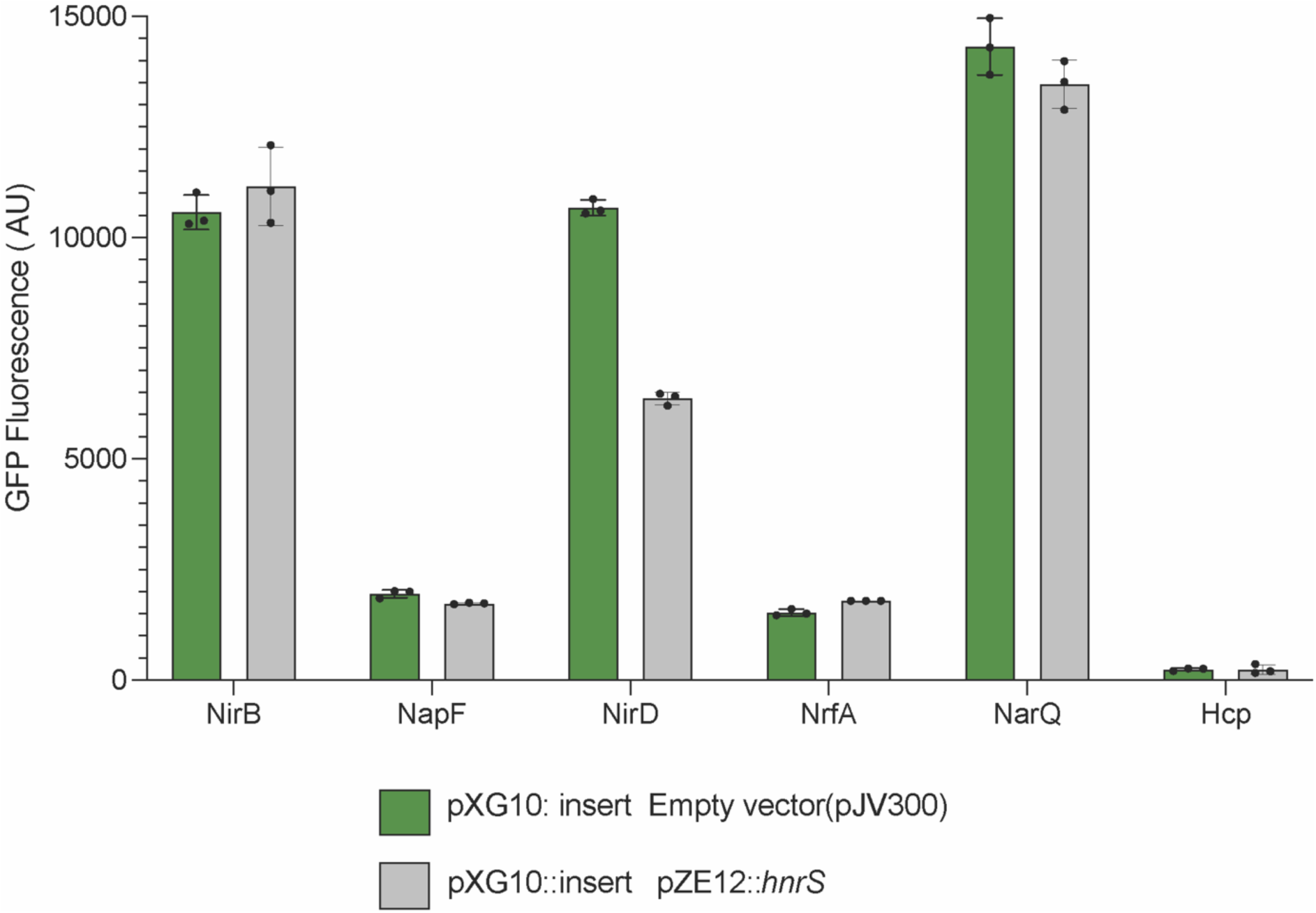
GFP translation fusion assay of genes involved in nitrate respiration. Bar graphs show fluorescence from translation fusions of NirB, NapF, NirD, NrfA, NarQ, and Hcp measured in the presence of HnrS or the control plasmid pJV300. The background control, pXG0, is included to account for autofluorescence, while insert (**-**) represents the empty **pXG10-SF** vector.

**Supplementary Figure 8.**
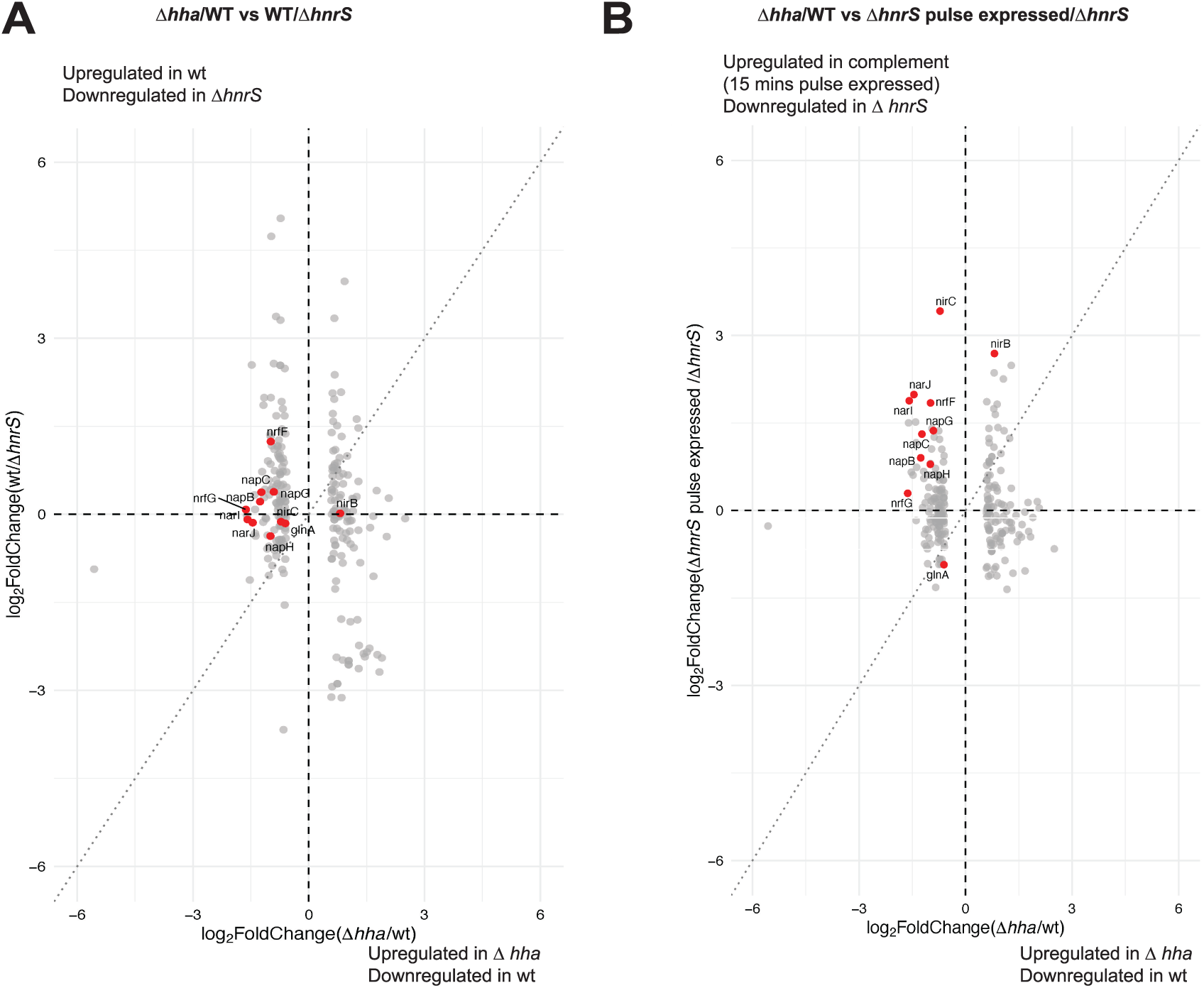
Comparative regulation of genes by Hha and HnrS. Scatterplot matrix showing relationships among Δ*hha*/WT and WT/Δ*hnrS*. Each panel represents pairwise comparisons of log₂ fold-change values from RNA-seq experiments. Red points indicate nitrogen metabolism genes, including *nirB, nirC, napG, napH, nrfF, nrfG, narI, narJ, napB, napC, glnA, gltB, gltD, ntrC,* and *amtB*, while grey points represent all other genes. Diagonal panels show density distributions of each dataset. Dashed lines indicate zero fold-change, and dotted lines indicate the diagonal (slope = 1) in pairwise scatter plots. Data are corrected such that the direction of regulation is consistent across comparisons (e.g., WT/Δ*hnrS* is inverted relative to Δ*hnrS*/WT).

## Notes

### Competing Interest Statement

The authors have declared no competing interest.

### Summary of Updates

The spelling was corrected for an author name.

