## Supplementary Methods for "Phage-encoded sRNA counteracts xenogenic silencing in pathogenic *E. coli*"

**Identification and analysis of HnrS within prophages**

The HnrS sequence was searched across multiple pathogenic *E. coli* strains using Nucleotide BLAST. Prophage regions were identified with PHASTER (1) and sequence alignments were performed to determine the conservation and genomic context of HnrS. Representative HnrS - encoding prophage regions were visualised using Easyfig, revealing that HnrS is consistently located upstream of key packaging and head protein genes. The analysis confirmed that HnrS is highly conserved within prophage regions and is restricted to these elements across all examined strains.

**Construction of EHEC deletion strains**

Chromosomal deletions of *hha* and *mazEFG* were performed using the pTOF25 allelic exchange system (2). For each target, ~500 nt regions flanking the gene(s) of interest were amplified from genomic DNA using primer pairs designed for Splicing by Overlap Extension (SOE) PCR:

***hha***: upstream flanking region, PP_hha_SmaI_A_R / PP_hha_SOE_B_F; downstream region, PP_hha_SOE_C_R / PP_hha_SalI_D_F.

***mazEFG***: upstream region, PP_mazE_SmaI_A_R / PP_mazE_SOE_B_F; downstream region, PP_mazF_SOE_C_R / PP_mazF_SalI_D_F.

Primers PP_hha_SOE_B_F / PP_hha_SOE_C_R (for *hha*) and PP_mazE_SOE_B_F / PP_mazF_SOE_C_R (for *mazEFG*) contained splicing-by-overlap-extension (SOE) to facilitate fusion of upstream and downstream fragments during SOE PCR. The resulting contiguous fragments were gel-purified and cloned into the pOF25 plasmid using SmaI and SalI restriction sites. Restriction digests were incubated at 37°C for 1 h, with FastAP thermosensitive alkaline phosphatase added after 45 min to prevent vector self-ligation, followed by enzyme inactivation at 65°C for 20 min. Ligations were performed with T4 DNA ligase at room temperature for 1 h and transformed into *E. coli* DH5α. Constructs were sequence-verified by Sanger sequencing.

A tetracycline resistance cassette (FRT–tetRA–FRT) was excised from pTOF1 using NotI and inserted into the pTOF25 allelic exchange plasmids. Transformants were selected on LB agar containing chloramphenicol (20 µg/mL) and tetracycline (10 µg/mL) at 30°C. Allelic exchange was promoted by serial passage on LB plates containing 6% sucrose (no NaCl) at 42°C. Recombinant colonies were screened for tetracycline resistance and chloramphenicol sensitivity, and deletion of the target gene (*hha* or *mazEFG*) was confirmed by colony PCR.

For *hha* and *mazEFG*, the tetRA cassette was subsequently excised by introducing pCP20, which encodes FLP recombinase, via electroporation. Correct deletion of each target was verified by PCR and sequencing.

**Construction and validation of chromosomal deletions and complementation plasmids**

Chromosomal deletions of *hha* and *mazEFG* in *E. coli* O157:H7 Sakai were validated through genomic DNA extraction using the Wizard® Genomic DNA extraction kit (Promega, Catalogue No. A1120) according to the manufacturer's instructions. Deletion of the targeted regions was confirmed by PCR amplification using gene-specific primers (PP_hha.del_F/PP_hha.del_R for *hha*; appropriate primers for *mazEFG*), and the resulting products were analyzed on a 1.5% agarose gel to detect the expected size reduction. Sanger sequencing was performed to further verify the deletions.

For complementation, medium copy number plasmids were used, with vector choice depending on the target gene. The *hha* gene, including 300 nucleotides upstream encompassing the native promoter and 103 nucleotides downstream of the stop codon, was amplified and cloned into pBR322. Insertion was confirmed using both plasmid-specific primers (pBR322_screening primer_F/R) and gene-specific primers (PP_hha.del_F/R), and Sanger sequencing verified the correct insert sequence.

Similarly, *mazEFG* complementation utilized pBR322 as the vector backbone. The *mazEFG* gene, including 143 nucleotides upstream (promoter) and 121 nucleotides downstream of the stop codon, was amplified using primers PP_comp_mazEFG_F1 and PP_comp_mazEFG_R1. EcoRI sites were incorporated into the primers, and the reverse primer included the rrnB terminator to ensure proper transcriptional termination. Insert integration was validated using pBR322-specific primers (pBR322_screening primer_F/R) as well as *mazEFG*-specific primers (PP_mazG_F and PP_mazE_R). Sequencing confirmed the correct construct.

For overexpression of *mazEFG*, the gene was cloned into the pZA21_MCS vector. The backbone was linearized using primers ZA21MCS.5P.R and ZA21MCS.HindIII.F, and the *mazEFG* insert was amplified with SL_mazE_KpnI_F and SL_mazG_HindIII_R. Both vector and insert were digested with HindIII, ligated, and transformed into competent cells. Transformants were screened by PCR and confirmed by sequencing.

**CRISPRi knockdown of sRNA HnrS**

The pCRISPathBrick (pCPB) vector was obtained from Addgene (USA) and used to perform targeted knockdown of HnrS in *E. coli* strain Sakai. Guide RNAs (sgRNAs) targeting hnrS were designed, 5′-phosphorylated, and annealed. The pCPB vector and annealed oligonucleotides were digested with BsaI and treated with FastAP to prevent self-ligation. Ligation of the digested vector with the sgRNA insert generated the pCPB::HnrS construct. Positive clones were confirmed by Sanger sequencing, and glycerol stocks were prepared for long-term storage. The verified pCPB::HnrS plasmid was then electroporated into *E. coli* Sakai stx(-) cells, which were selected on LB agar containing kanamycin and chloramphenicol.

**RNA extraction and Northern Blot**

RNA was extracted using the guanidinium thiocyanate–phenol method (3) with a few modifications. Bacterial cultures were grown in LB broth, minimal M9 medium, or MEM-HEPES supplemented with 0.1% glucose and 250 nM $Fe(NO_3)_3$ until they reached the desired growth phase, as determined by $OD_{600}$. Once the cultures reached the appropriate density, cells were collected by centrifugation at 4000 × g for 10 minutes at 4°C. The supernatant was discarded, and the cell pellet was resuspended in 1 mL of a 1:1 mixture of GTC buffer and phenol (GTC buffer composition: 4M guanidinium thiocyanate, 2% Sarkosyl, 50 mM Tris pH 8.0, 10 mM EDTA, 1% β-mercaptoethanol), along with 250 μL of 0.1 mm zirconia-silica beads

(Biospec products, Catalogue No. 11079101z). Cells were lysed by vortexing and then incubated in a 65°C water bath for 5 minutes. The lysate was transferred to a fresh tube containing 350 µL of chloroform and 120 µL of 3M NaOAc (pH 5.2), vortexed for 20 seconds, and centrifuged at maximum speed for 5 minutes at room temperature to separate the phases. The aqueous phase was carefully transferred to a new tube containing 550 µL of phenol:chloroform:isoamyl alcohol (25:24:1), vortexed, and centrifuged again as before. The aqueous phase was then transferred to 450 µL of chloroform and processed in the same manner. The final aqueous phase was transferred to two and a half volumes of 100% ethanol and 15 µg/mL Glycoblue coprecipitant (ThermoFisher, Catalogue No. AM9515) and incubated at -80°C overnight for ethanol precipitation. The following day, the samples were centrifuged at 16,000 × g for 30 minutes at 4°C. The ethanol was aspirated, and the RNA pellet was washed twice with 700 µL of 70% ethanol, followed by centrifugation at 16,000 × g for 20 minutes at 4°C. After the final spin, the remaining ethanol was aspirated, and the pellet was left to air dry before being resuspended in 40 µL of nuclease-free water.

Electrophoresis was performed on 8% polyacrylamide gels in TBE-urea buffer (1x TBE, 8M urea, 8% acrylamide). Prior to casting the gel, TEMED (tetramethylethylenediamine) and 1% APS (ammonium persulfate) were added as polymerization initiators. The gel was pre-warmed by running it for 30 minutes at 200V to ensure even heating. RNA samples were prepared by mixing them with 2x formamide loading buffer (80% formamide, 10 mM EDTA, 0.025% bromophenol blue, 0.025% xylene cyanol) and incubated at 65°C for 10 minutes to denature any secondary structures. The denatured samples were loaded onto the gel and electrophoresed at 200V for at least 3 hours in a gel measuring 16.5 x 14.5 x 0.4 mm.

Polyacrylamide gels containing the separated RNA was transferred to a nylon membrane using a CBS Scientific Large Blotting System (Catalogue No. EBU-102). The transfer was performed at 30V in 0.5X TBE buffer overnight. The RNA was then immobilised on the membrane by UV-crosslinking using a Stratagene Auto-Crosslinker, with an exposure of 1200 mJ of UV light. After immobilization, the membrane was prehybridised by incubating it in Ambion™ ULTRAhyb™ Ultrasensitive Hybridization Buffer (Catalogue No. AM8670) at 30°C for 30 minutes to block non-specific binding sites.

Oligonucleotide probes (30-35 nucleotides long) were designed using NetPrimer and Integrated Genome Browser. These probes were labelled with 32P-ATP using T4 polynucleotide kinase (PNK). The labelling reaction consisted of 1.5 μL of 10 mM oligonucleotide probe, 2.5 μL of 32P-ATP, 1.5 μL of 10X PNK buffer, 1 μL of T4 PNK, and 9 μL of MilliQ water. The reaction was incubated at 37°C for 1 hour. The radiolabelled probes were purified using a GE Healthcare Illustra Microspin™ G-50 column (Catalogue No. 27-53301) to remove excess nucleotides, following the manufacturer's instructions.

The purified radiolabelled probes for HnrS detection BS_HnrS_NB (Table 3.4) were then added to the prehybridised membrane and incubated overnight at 30°C in a humidified chamber to allow for optimal hybridization. The next day, the membrane was washed three times by incubating it with 2x SSPE buffer (0.3M NaCl, 20 mM NaH2PO4, 2 mM EDTA) containing 0.1% SDS for 10 minutes at 42°C to remove any unbound probe. The membrane was then exposed to a BAS Storage Phosphor Screen and imaged using a Typhoon™ FLA9500 imaging system to detect the radiolabelled signal.

**HnrS prevalence assay in *E. coli* genomes**

Genomic data for 3,230 E. coli strains were retrieved from the NCBI database and processed using a series of bioinformatics tools to identify key genetic markers and assess the strains' pathogenic potential. Initially, the data underwent K-mer Analysis (KMA) to detect essential housekeeping genes such as *recA*, *purA, mdh, icd, gyrB, fumC,* and *adk,* which were used to confirm strain identity and estimate genome depth. In addition, KMA identified pathotyping genes like *eae, escV, stx1,* and *stx2,* allowing for the classification of strains into distinct E. coli pathovars, including EHEC, EPEC, EAEC, and others. A detailed list of these pathotyping genes, along with their associated pathovars, is provided in Table A

To determine the copy number of HnrS, a relative coverage approach was used based on the KMA results. First, the sequencing reads for each strain were processed using the test_commandPP script (Section 3.3.5), which runs KMA along with ShigeFinder and STECFinder to identify genes present in the sample. During this process, the KMA output (*_kma_out.res) was analysed to identify the coverage depth of the HnrS gene among other genes. The HnrS gene's coverage was extracted from the KMA results, and the relative coverage was calculated by comparing the observed coverage of HnrS to the overall genomic depth of the strain, using housekeeping genes for normalisation. This relative coverage of HnrS was then used as an indicator of its copy number, with higher relative coverage suggesting a greater number of copies of the HnrS gene present in the strain. This calculation was done in the post-processing step through the Python script process_ecolitypingv3, which interpreted the KMA results and calculated the relative coverage for each strain.

Table A. Pathotying genes with associated pathovars.

| S.N | Gene | Pathovar | Reference |
|---|---|---|---|
| **1.** | *fyuA* | UPEC | (4) |
| **2.** | *papC* | UPEC | (4) |
| **3.** | *hlyA* | UPEC | (4) |
| **4.** | *traT* | UPEC | (4) |
| **5.** | *escV* | LEE positive strains (EPEC, ATEC, STEC) | (5) |
| **6.** | *bfp* | EPEC, atypical, Samonella | (5) |
| **7.** | *stx1* | STEC | (5) |
| **8.** | *stx2* | STEC | (5) |
| **9.** | *aaic* | EAEC | (5) |
| **10.** | *aggR* | EAEC | (5) |
| **11.** | *invA* | EIEC | (5) |
| **12.** | *uidA* | E.coli | (5) |
| **13.** | *eae* | EPEC and EHEC | (5) |
| **14.** | *ipaH* | EIEC and Shigella | (6) |
| **15.** | *daaD* | DAEC | (5) |
| **16.** | *virG* | EIEC and Shigella | (6) |
| **17.** | *est* | ETEC | (5) |
| **18.** | *elt* | ETEC | (5) |

Shigella Finder was used to categorise strains as Shigella, Enteroinvasive E. coli (EIEC), or non-Shiga toxigenic E. coli. Based on specific genetic markers, strains were further classified into "Shigella," "EIEC," or "Shigella/EIEC Unclustered," with additional sub-classifications based on serotype. Meanwhile, STEC Finder was used to identify serotypes within the STEC group by detecting serotype-specific markers such as *stx1*, *stx2*, *eae*, and *escV*. This enabled precise classification of strains into known serotypes, including O157:H7, and facilitated grouping them into categories like EHEC, based on their genetic profile. The STEC Finder software relies on a predefined database of serotype-specific markers to assign each strain to its corresponding serotype.

To further refine pathovar classification, a Boolean logic expressions were applied, as detailed in Table B. This algorithm enabled classification of strains into specific pathovars based on the presence or absence of key genes. For instance, EHEC strains were identified by the presence of *eae* or *escV* along with *stx1* or *stx2* but without *bfp*, while Uropathogenic Escherichia coli (UPEC) strains were defined by multiple virulence factors such as *fyuA*, *traT*, *hlyA*, and *papC*.

Table B. Boolean logic expression to classify pathovars

| *Pathovar* | *Boolean Logic Expression* |
|---|---|
| *EHEC* | ((*eae* OR *escV*) AND *stx*) NOT *bfp* |
| *EPEC* | (*eae* OR *escV)* NOT *stx* |
| *EAEC* | (*aggR* AND *aaiC*) NOT *stx* |
| *DAEC* | (*daaD* OR *afaD*) NOT (*stx* OR *bfp*) |
| *UPEC* | ((*fyuA* AND *traT*) OR (*fyuA* AND *hlyA*) OR (*fyuA* AND *papC*) OR (*traT* AND *hlyA*) OR (*traT* AND *papC*) OR (*hlyA* AND *papC*)) |
| *ETEC* | (*elt* OR *est*) |
| *EIEC* | (*invA* AND *ipaH*) AND NOT (*stx* OR *bfp*) |
| *Shigella* | (*ipaH* and *virG*) |
| *Salmonella* | (*invA*) |

Finally, the relative coverage of the HnrS gene was assessed by comparing its coverage to the overall genome depth, providing a measure of its prevalence across different E. coli strains. The final output compiled essential information for each strain, including strain ID, Shigella/EIEC status (from Shigella Finder), serotype (from STEC Finder), gene-based pathotype (from KMA), and the list of detected pathotyping genes. This dataset was used to assess the occurrence of HnrS across various genomes and strains, aiding in the prediction of its presence in virulent strains.

Codes for performing HnrS prevalence analysis in *E. coli* genomes is in https://github.com/pranitapoudyal/EcOnc10-prevalence-in-E.-coli-genomes-with-M.-Payne

**Construction of HnrS deletion in EHEC str. Sakai using CRISPR-Cas9**

CRISPR-Cas9 was used to genetically modify *E. coli* O157:H7 strain Sakai stx+. The process involved a two-plasmid system as described in (7), where sgRNA was inserted into pTarget-F using inverse PCR with Phusion using BS_EcOnc10_sgRNA_F and BS_pTargetF_R_5P (5' phosphorylated). After amplification, 1ul of DpnI enzyme was added to the reaction and incubated for 1 hour at 37°C, followed by purification of the linearised vector using the Wizard® SV Gel and PCR Clean-Up System (Promega, Catalogue No. A9282). The purified amplicon was then ligated using T4 DNA ligase (ThermoFisher) and transformed into competent DH5α cells via heat shock. Plasmids were extracted and confirmation of the insertion of HnrS into pTargetF plasmid was confirmed through Sanger sequencing using BS_pTarget_F screening primer and the plasmid was termed as pTargetF::HnrS. Flanking regions for homologous recombination ~300 nucleotides each extending 5' and 3' of HnrS were amplified from *E. coli* O157:H7 strain Sakai stx+ using the primer pairs BS_EcOnc10_Down_F, BS_EcOnc10_Down_R and BS_EcOnc10_UP_F, BS_EcOnc10_UP_R primers. The two arms were spliced by Gene Splicing by Overlap Extension (SOEing) using the primers BS_EcOnc10_Down_R and BS_EcOnc10_UP_F generating a single product that excludes HnrS.

The plasmid pTargetF::HnrS and the homology arms were double digested with *HindIII-HF* and *XhoI*(New England Biolabs, USA, Catalog R3104S R0146S). Plasmid pTargetF::HnrS was incubated with 1 unit of Fast AP thermosensitive alkaline phosphatase (ThermoFisher Scientific, USA, Catalog EF0651) at 45 mins in order to decrease self ligation and the reactions were inactivated by incubating at 65°C for 20 mins. A 20µl ligation reaction mix (1X T4 DNA ligase buffer and 1 unit of T4 ligase (ThermoFisher Scientific, USA, EL0011) was prepared with a 1:3 molar ratio of vector and insert at 20°C, for 1.5 hours. The plasmid containing both the sgRNA and the repair template for homologous recombination are termed pTargetT::HnrS.

The plasmid pTargetT::HnrS was transformed into DH5α competent cells using heat shock transformation. Selection of the colonies were performed on LB plates supplemented with spectinomycin and screened for homology arm insertion using primers BS_EcOnc10_Down_F and BS_EcOnc10_UP_R. Plasmids were extracted and confirmation of the insertion of homology arms was confirmed through Sanger sequencing using BS_EcOnc10_Down_F primers.

All incubations occurred at 30°C unless specified otherwise. The pCas plasmid was electroporated into *E. coli* O157:H7 strain Sakai stx+. Transformants were streaked out for single colonies on LB-Km50 plates. Single colonies were incubated overnight in LB-Km50 broth, then subcultured 1/100 the next day. Once an $OD_{600}$ of 0.45-0.6 was reached, λ-Red was induced by adding 50 mM of L-arabinose for 30 minutes. Cells were harvested and the desired pTargetT plasmid was electroporated as above. Transformants were plated onto LB-Km50Sm50 plates. Colony PCR was done to verify that the desired modification was present, and plasmids were cured by streaking out positive colonies onto LB-Km50 containing 0.5 mM IPTG and incubated at 42°C overnight.

**Complementation of an HnrS complementation plasmid**

Inverse PCR was performed to clone HnrS into pBAD+1 vector. The vector pBAD+1 was cloned with 66 nucleotides of HnrS using the following primers pBAB_PP_EcOnc10_F and pBAD_PP_5P_R. The amplicons were digested with 1µl of DpnI(NEB) and incubated for 1 hour at 37°C cleaned up using the Promega PCR cleanup kit, ligated overnight at 16°C and transformed into DH5a then into EHEC str Sakai.

**Sanger sequencing of mutants**

Genomic DNA was extracted using the Wizard® Genomic DNA Extraction Kit (Promega, Catalogue No. A1120) to verify successful chromosomal deletion and repair. Primers flanking the chromosomal region of interest were designed and used to amplify the target sequence by PCR. The PCR product was then purified using the Wizard® SV Gel and PCR Clean-Up System (Promega, Catalogue No. A9282). The purified PCR product was sent for Sanger sequencing with one of the primers. The resulting sequence, obtained in FASTA format, was analysed by aligning it with the original reference sequence in Benchling to confirm the presence of the chromosomal changes.

**Phage plaquing assay**

The double-agar overlay technique was used to measure plaque formation and determine phage titers (8, 9). Square plastic petri dishes measuring 10 cm × 10 cm were filled with approximately 35 mL of LB agar as the bottom layer, consisting of 10 g/L tryptone, 5 g/L yeast extract, 10 g/L NaCl, and 2% agar, and allowed to dry. The top agar layer, consisting of 10 g/L tryptone, 5 g/L yeast extract, 10 g/L NaCl, and 0.5% agar, was maintained at 65°C in a water bath. Once cooled to room temperature, 10 mL of the top LB agar was supplemented with 10 mM $CaCl_2$, 10 mM $MgSO_4$, and 800 µL of indicator bacteria (overnight cultures of *E. coli* MG1655 with or without EcOnc10). For Stx phage assays, 1.5 µg/mL mitomycin C (MMC) was incorporated into the top agar. After briefly vortexing, the top agar mixture was poured over the dried bottom agar layer and allowed to set. Serial dilutions of purified phages were prepared using LB media with 5 mM $CaCl_2$, and 5 µL of each dilution was spot plated onto the top agar. The plates were incubated overnight at 37°C, and plaques were enumerated the following day.

**Knock-down of HnrS using CRISPRi in EHEC strain Sakai**

To understand the function of HnrS and to study the effects of reduced gene expression on cellular pathways and phenotypes gene knockdown was performed.

**Generation of the CRISPRi gene silencing system**

A modular system for multigene repression in E. coli using CRISPRi (10) has been developed. This system, named CRISPathBrick, enables the insertion of up to five target guide RNAs into a single plasmid, pCPB, through Spacer Repeat Bricks (SRBs). The plasmid pCPB also encodes dCas9, a modified Cas9 protein that lacks nuclease activity, which effectively attenuates gene expression and generates knockdowns.

**CRISPRi knockdown of sRNA HnrS**

The pCRISPathBrick (pCPB) vector was obtained from Addgene (USA). SRB oligo nucleotides that were 5' phosphorylated containing the sgRNA (specified in Table 2.3) were designed and ordered from IDTDNA. Annealing of the SRB oligonucleotides involved combining them in equimolar (50 µM total) amounts and performing the annealing process in an IDTDNA duplex buffer(100 mM potassium acetate; 30 mM HEPES, pH 7.5). Gel electrophoresis confirmed successful annealing.

The pCPB vector (500 ng) and annealed oligonucleotides (200 ng) were digested using BsaI enzyme, with FastAP thermosensitive alkaline phosphatase added to prevent self-ligation. BsaI was then inactivated. Ligation of the pCPB vector (40 ng) and the annealed oligonucleotide insert (80 ng) occurred at room temperature (21°C) for 1.5 hours, resulting in the formation of the desired construct, pCPB::HnrS. Positive colonies were subjected to Sanger sequencing to verify the presence of the desired construct. A glycerol stock was prepared from a colony confirmed to carry the correct sequence for long-term storage, and the extracted pCPB::HnrS plasmid was electroporated into E. coli str. Sakai stx(-) cells, which were plated on LB agar supplemented with kanamycin and chloramphenicol.

### Construction of GFP translational fusions and sRNA expression vectors

GFP translational fusions were constructed following the protocol for operon fusion (11) as outlined in (12) where primers were designed to amplify the DNA fragment from a polycistronic locus, rather than a single gene for fusion with *lacZ* and GFP. The sense primer was designed to anneal to the C-terminal coding region of the upstream gene within the operon, including the interaction site (seed sequence), spanning 87 base pairs that encode the last 29 amino acids of the *mazF* gene. This primer also included an NsiI restriction site extension at the 5' end (sequence: ATGCAT), ensuring that the restriction site was in frame with the upstream gene. The antisense primer annealed to the N-terminal coding region of the downstream *mazG* gene within the locus, spanning 84 base pairs encoding first 27 amino acids. This primer carried an NheI site extension (sequence: GCTAGC), which encodes the second and third residues of GFP. The NheI site was also positioned in frame with the downstream gene. Careful checks were made to ensure that the amplified sequence did not contain any internal NsiI or NheI sites, which would need to be avoided. The resulting product was cloned into the pXG30SF vector, which encodes superfolder GFP under the control of the PLtetO-1 promoter, using NheI and NsiI (FastDigest, ThermoFisher).

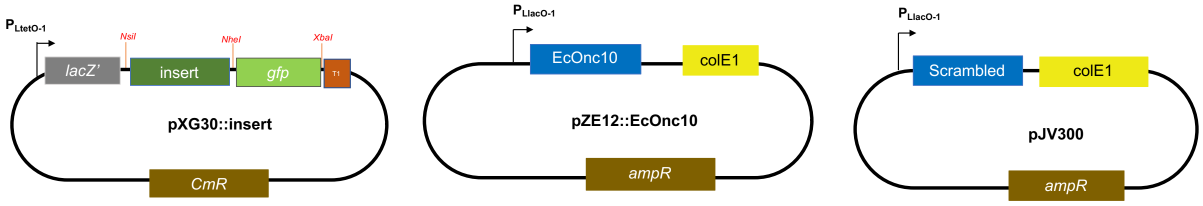

HnrS was cloned into pZE12 by inverse PCR with Phusion using pZE12-inverse PCR.F and pZE12-inverse PCR.5P.R (5’ phosphorylated) where pZE12-inverse PCR.F contains the sequence for HnrS. Following amplification, it was purified using the Wizard ® SV Gel and PCR Clean-Up System, the purified amplicon was ligated using T4 DNA ligase (ThermoFisher) reaction to circularise the vector. The reaction was incubated at room temperature for 1 hour before heat-shock transformation into ultra-competent DH5α cells. Colonies obtained were screened using screening primers pZE12.seq.F and PZE12.seq.R Successful insertion was checked for the appropriate band size and were chosen. Plasmids were extracted using the Wizard® Plus SV Minipreps DNA Purification System (Promega) according to the manufacture's protocol, and insertion of HnrS was confirmed via Sanger sequencing using F screening primers (Supplementary Table 1).

**Determination of bacterial motility**

Bacterial strains were streaked on solid LB-agar plates and incubated at 37°C overnight. From these plates, a single colony was subcultured in LB broth containing ampicillin (100 μg/mL) until it reached an OD600 of 0.6. 0.2μL of culture was patched onto an LB Agar plate with a lower agar concentration (0.2%) and incubated at 30°C. Bacterial motility was observed, and diameter of the flare was noted.

For single cell motility assays, overnight bacterial cultures were subcultured at a 1:100 dilution in Tryptone Broth (1% (w/v) tryptone, 0.5% (w/v) NaCl) to achieve an OD600 of 0.1. The cultures were allowed to grow to the late-exponential growth phase. To prepare the samples for imaging, the cell cultures were further diluted to approximately 1 × 105 cells/mL. The diluted bacterial suspension was added into a flow cell, which consisted of a microscope slide and a cover slip separated by tape, creating a confined tunnel like space for the bacteria (13). Before imaging, the prepared sample was placed on the microscope stage for 10 minutes to reach thermal equilibrium.

Bacterial motility was observed using a Nikon phase-contrast microscope with a 40× objective lens, and time-lapse videos were recorded using a Chameleon3 CM3 camera (Point Grey Research), capturing 20-second videos at 20 frames per second for a total of 400 frames.

Swimming speeds were estimated using a custom LabVIEW 2023 software (14) and subsequently plotted against bacterial strains using GraphPad Prism 8.

**Type III secretion assay of secreted proteins**

Type III secretion (T3S) assay was performed as previously described by (15). Overnight LB broth cultures were diluted 1:100 in MEM-HEPES (supplemented with 250nM of $Fe(NO_3)_3$ and 0.1% glucose). Cultures were grown to an $OD_{600}$ of 0.8 and supernatants were collected by centrifugation at 4000 rpm for 30 min. Supernatants were syringe filtered through 0.45µm low protein-binding filters (Millex-HP Syringe Filter Unit) and secreted proteins were precipitated overnight at 4°C with 10% trichloroacetic acid (TCA; Sigma-Aldrich) and 100µg of rAlbumin (Recombinant Albumin; New England BioLabs, Catalogue No. B9200S) was added as a co-precipitant and loading control. Secreted proteins were centrifuged at 4000g for 30 mins at 4°C. Pellets were air-dried, resuspended in 150µl of loading dye and heated for 10 mins at 95°C and analysed through a SDS-PAGE ( 4-20% Mini- PROTEAN TGX PreCast Gels Catalogue No. 4561093). SDS-PAGE gels were stained with Commassie and imaged on a Bio-Rad ChemiDoc using default white light illumination setting.

Proteins separated by SDS-PAGE were transferred onto a Protran Nitrocellulose membrane (Cytiva Amersham, Catalogue No. 15249794) using a Trans-Blot electrophoretic transfer cell (Bio-Rad). The nitrocellulose membrane was blocked with 8% (w/v) dried milk powder in phosphate-buffered saline (PBS) at room temperature for 1 hour, followed by three washes with PBS.

Immunoblotting was performed to detect EspD proteins. Mouse monoclonal anti-EspD antibodies (kindly provided by Professor T. Chakraborty, University of Giessen) and rabbit polyclonal anti-mouse IgG HRP-conjugated secondary antibodies (Cell Signaling Technology, Catalogue No. 7076) were both diluted 1:2000. The membrane was incubated with the primary antibody overnight at 4°C on a platform shaker and subsequently washed three times for 10 minutes in PBS. The secondary antibody incubation was performed for 1 hour at 4°C on a platform shaker, followed by three 10-minute washes in PBS.

For enhanced chemiluminescence (ECL) detection, the membranes were incubated in 2.5 mL of ECL Solution 1 (Amersham) mixed with 2.5 mL of ECL Solution 2 (Amersham) for 5 minutes at room temperature. Chemiluminescence was detected by imaging the nitrocellulose membrane using the ChemiLITE Chemiluminescence Imaging System (Cleaver Scientific).

As a control for protein loading, mouse monoclonal anti-DnaK antibody (Roslin Institute, University of Edinburgh) was used as a housekeeping gene marker in the Western blot, which was performed on whole-cell lysates from the same secreted protein sample. The same procedure as outlined above was followed, using the same dilution for the primary antibodies, followed by incubation with rabbit polyclonal anti-mouse IgG HRP-conjugated secondary antibodies.

**LEE1, LEE4 and LEE5 translation GFP reporter assays.**

Plasmid-based LEE1, LEE4, and LEE5 promoter and 5'UTR fusions to green fluorescent protein (GFP) were acquired from Prof David Gally (16). The native promoters and corresponding coding sequences for key genes in each operon —*ler* for LEE1, *tir* for LEE5, and *sepL* for LEE4 —cloned into the pAJR70 vector to create translational fusions for each operon. All constructs were verified by sequencing to ensure correct insertion and integrity (16).

The GFP fusion plasmids were then introduced into EHEC strain Sakai pBR322 (wild-type), EHEC strain Sakai Δ*hha* pBR322 (*hha* deletion), and EHEC strain Sakai Δ*hha* pBR322::*hha* (complemented) by electroporation (Section 2.2.7). Transformants were selected on LB agar plates containing 100 µg/mL ampicillin and 50 µg/mL chloramphenicol. Bacterial cultures were grown in MEM-HEPES medium supplemented with chloramphenicol, ampicillin, glucose, and iron to promote bacterial growth and reporter expression. $OD_{600}$ and fluorescence were measured every hour in a black 96-well plate (Fluoro-Nunc) using a fluorimeter (FLUOstar Optima). $OD_{600}$ was monitored to track growth, while fluorescence was measured to quantify GFP expression. Fluorescence data were plotted against $OD_{600}$ values using GraphPad Prism software. The promoter-less plasmid pAJR70 served as a negative control to assess any background fluorescence from the strain or medium.

**Adhesion Assay**

Coverslips were placed in 24-well plates and coated with 200 µL of collagen overnight at 4°C. The following day, wells were gently washed with PBS to remove excess collagen and allowed to air-dry for 3–4 hours. Bovine epithelial cells (EBLs) were seeded at approximately $1 \times 10^5$ cells per well and incubated overnight at 37°C with 5% $CO_2$ to allow adherence. Overnight bacterial cultures were sub-cultured 1:10 in MEM supplemented with 0.1% glucose and 250 nM $Fe(NO_3)_3$ and incubated for approximately 3 hours. One hour prior to infection, the EBL growth medium was replaced with MEM to match the bacterial growth medium, and cells were then infected with 100 µL of bacterial culture to achieve a multiplicity of infection (MOI) of ~100. Infected cultures were incubated for 3 hours at 37°C, after which non-adherent bacteria were removed by two washes with MEM. Following an additional 2-hour incubation, wells were washed twice with phosphate-buffered saline (PBS), and cells were fixed with 100 µL of 1% paraformaldehyde (PFA) per well for 10 minutes at room temperature. Fixed cells were washed with PBS and stored at 4°C until immunostaining.

For immunostaining, fixed cells were permeabilized with 200 µL of 0.1% Triton X-100 per well for 10 minutes at room temperature and washed twice with PBS. Cells were blocked with 200 µL of 1 mg/mL bovine serum albumin (BSA) in PBS for 1 hour at room temperature and washed twice with PBS. Primary staining was performed using 200 µL of rabbit anti-O157 antibody (1:50 dilution) for 1 hour at room temperature, followed by three 5-minute washes with PBS. Secondary staining was carried out with 200 µL of goat anti-rabbit IgG Alexa Fluor 568 (1:1000 dilution) for 1 hour at room temperature in the dark, followed by three 5-minute washes with PBS. Actin filaments were stained using 200 µL of Phalloidin Alexa Fluor 488 (BioLegend) for 30 minutes at room temperature in the dark, followed by two washes with PBS, and nuclei were stained with 100 µL of Hoechst 33342 (1:2000 dilution) for 5–10 minutes at room temperature in the dark, followed by three washes with PBS.

Coverslips were then carefully removed and inverted onto a drop of ProLong antifade mounting medium on glass slides and allowed to cure for 24 hours. Imaging was performed at at the Katherina Gaus Centre for Microscopy (UNSW) using a Zeiss LSM900 inverted confocal laser scanning microscope with a 63× oil immersion objective. Fluorescence was detected using 488 nm excitation for Phalloidin, 405 nm for Hoechst, and 633 nm for Alexa Fluor 568, and images were processed and merged using ImageJ software.

**Gene Ontology (GO) enrichment analysis**

To investigate the functional roles of differentially expressed genes between the EcOnc10 deletion mutant and the wild-type strain, Gene Ontology (GO) enrichment analysis was performed in R using the clusterProfiler package. Genes with an absolute log2 fold change ≥ 1, as determined by DESeq2, were included in the analysis. The enrichment was assessed across the three GO categories: biological processes, molecular functions, and cellular components, using the most recent GO annotation available for *E. coli* O157:H7 str. Sakai. Enriched terms were visualised as scatter plots generated in ggplot2, with the size of each point representing the number of genes associated with the term and colour indicating the adjusted p-value significance. Figure of gene ontology enrichment analysis is in Supplementary figure 2.


1. Arndt D, Grant JR, Marcu A, Sajed T, Pon A, Liang Y, Wishart DS. 2016. PHASTER: a better, faster version of the PHAST phage search tool. Nucleic Acids Res 44:W16-21.
2. Merlin C, McAteer S, Masters M. 2002. Tools for characterization of Escherichia coli genes of unknown function. J Bacteriol 184:4573-81.
3. Tollervey D, Mattaj IW. 1987. Fungal small nuclear ribonucleoproteins share properties with plant and vertebrate U-snRNPs. Embo j 6:469-76.
4. Kudinha T, Kong F, Johnson JR, Andrew SD, Anderson P, Gilbert GL. 2012. Multiplex PCR-Based Reverse Line Blot Assay for Simultaneous Detection of 22 Virulence Genes in Uropathogenic Escherichia coli. Applied and Environmental Microbiology 78:1198-1202.

5. Müller D, Greune L, Heusipp G, Karch H, Fruth A, Tschäpe H, Schmidt MA. 2007. Identification of unconventional intestinal pathogenic Escherichia coli isolates expressing intermediate virulence factor profiles by using a novel single-step multiplex PCR. Appl Environ Microbiol 73:3380-90.
6. Vidal M, Kruger E, Durán C, Lagos R, Levine M, Prado V, Toro C, Vidal R. 2005. Single multiplex PCR assay to identify simultaneously the six categories of diarrheagenic Escherichia coli associated with enteric infections. J Clin Microbiol 43:5362-5.
7. Jiang Y, Chen B, Duan C, Sun B, Yang J, Yang S. 2015. Multigene editing in the Escherichia coli genome via the CRISPR-Cas9 system. Appl Environ Microbiol 81:2506-14.
8. Islam MR, Ogura Y, Asadulghani M, Ooka T, Murase K, Gotoh Y, Hayashi T. 2012. A sensitive and simple plaque formation method for the Stx2 phage of Escherichia coli O157:H7, which does not form plaques in the standard plating procedure. Plasmid 67:227-35.
9. Bonanno L, Loukiadis E, Mariani-Kurkdjian P, Oswald E, Garnier L, Michel V, Auvray F, Griffiths MW. 2015. Diversity of Shiga Toxin-Producing Escherichia coli (STEC) O26:H11 Strains Examined via <i>stx</i> Subtypes and Insertion Sites of Stx and EspK Bacteriophages. Applied and Environmental Microbiology 81:3712-3721.
10. Cress BF, Toparlak Ö D, Guleria S, Lebovich M, Stieglitz JT, Englaender JA, Jones JA, Linhardt RJ, Koffas MA. 2015. CRISPathBrick: Modular Combinatorial Assembly of Type II-A CRISPR Arrays for dCas9-Mediated Multiplex Transcriptional Repression in E. coli. ACS Synth Biol 4:987-1000.
11. Menz L. 2017. The role of the small RNA ecOnc10 in regulating MazEF in enterohaemorrhagic Escherichia coli O157:H7. Bachelor of Biotechnology. UNSW, Sydney.
12. Urban JH, Vogel J. 2007. Translational control and target recognition by Escherichia coli small RNAs in vivo. Nucleic Acids Res 35:1018-37.
13. Palma V, Gutiérrez MS, Vargas O, Parthasarathy R, Navarrete P. 2022. Methods to Evaluate Bacterial Motility and Its Role in Bacterial-Host Interactions. Microorganisms 10.
14. Ishida T, Ito R, Clark J, Matzke NJ, Sowa Y, Baker MAB. 2019. Sodium-powered stators of the bacterial flagellar motor can generate torque in the presence of phenamil with mutations near the peptidoglycan-binding region. Mol Microbiol 111:1689-1699.
15. Roe AJ, Yull H, Naylor SW, Woodward MJ, Smith DG, Gally DL. 2003. Heterogeneous surface expression of EspA translocon filaments by Escherichia coli O157:H7 is controlled at the posttranscriptional level. Infect Immun 71:5900-9.
16. Fernandez-Brando RJ, Yamaguchi N, Tahoun A, McAteer SP, Gillespie T, Wang D, Argyle SA, Palermo MS, Gally DL. 2016. Type III Secretion-Dependent Sensitivity of Escherichia coli O157 to Specific Ketolides. Antimicrob Agents Chemother 60:459-70.
